# Precocious Chondrocyte Differentiation Disrupts Skeletal Growth in Kabuki Syndrome Mice

**DOI:** 10.1101/599878

**Authors:** Jill A. Fahrner, Wan-Ying Lin, Ryan C. Riddle, Leandros Boukas, Valerie B. DeLeon, Sheetal Chopra, Susan E. Lad, Teresa Romeo Luperchio, Kasper D. Hansen, Hans T. Bjornsson

**Author notes:** Authorship notes: S.E.L.’s new affiliation is the Department of Biological Sciences, University of Notre Dame, Notre Dame, IN. S.C.’s new affiliation is College of Osteopathic Medicine, Kansas City University of Medicine and Biosciences, Kansas City, MO. Conflict of interest statement: H.T.B. is a consultant with Millennium Therap.

## Abstract

Kabuki syndrome 1 (KS1) is a Mendelian disorder of the epigenetic machinery caused by mutations in the gene encoding KMT2D, which methylates lysine 4 on histone H3 (H3K4). KS1 is characterized by intellectual disability, postnatal growth retardation, and distinct craniofacial dysmorphisms. A mouse model (*Kmt2d*^*+/ β Geo*^) exhibits features of the human disorder and has provided insight into other phenotypes; however, the mechanistic basis of skeletal abnormalities and growth retardation remains elusive. Using high-resolution micro-computed tomography we show that *Kmt2d*^*+/βGeo*^ mice have shortened long bones and ventral bowing of skulls. *In vivo* expansion of growth plates within both the skull and long bones suggests disrupted endochondral ossification as a common disease mechanism. Stable chondrocyte cell lines harboring inactivating mutations in *Kmt2d* exhibit increased proliferation and differentiation, which further supports this mechanism. A known inducer of chondrogenesis, SOX9, and its targets show markedly increased expression in *Kmt2d*^*-/-*^ chondrocytes. By transcriptome profiling, we identify *Shox2* as a putative KMT2D target. We propose that decreased KMT2D-mediated H3K4me3 at *Shox2* releases *Sox9* inhibition and thereby leads to enhanced chondrogenesis, providing a novel and plausible explanation for precocious chondrocyte differentiation. Our findings not only provide insight into the pathogenesis of growth retardation in KS1, but also suggest novel therapeutic targets to rescue growth retardation in KS1 and related disorders.

## Introduction

Mendelian disorders of the epigenetic machinery disrupt the fundamental processes of neurological development and growth (1). This rapidly growing group of inherited conditions resulting from germ-line mutations in components of the epigenetic machinery is expected to have broad epigenomic consequences. Despite growth abnormalities being the second most common disease manifestation, molecular underpinnings have not been examined in detail but could provide insight into disease mechanisms that may be broadly applicable to other more common growth disorders, like idiopathic short stature.

Kabuki syndrome (KS; MIM 147920), an autosomal dominant Mendelian disorder of the epigenetic machinery (1), results from heterozygous typically *de novo* inactivating mutations in *KMT2D* (2), which encodes an epigenetic writer that normally catalyzes histone methylation on H3 at lysine 4 (H3K4me). Individuals with Kabuki syndrome 1 (KS1) exhibit the cardinal phenotypic features of postnatal growth retardation, intellectual disability, and craniofacial abnormalities (3,4). The distinct craniofacial features that are characteristic of KS1 include flattening of the facial profile, elongated palpebral fissures with eversion of the lower eyelids, highly arched and interrupted eyebrows, short columella with a depressed nasal tip, prominent ears, and palate abnormalities; these often provide the best clue for clinical diagnosis in affected individuals (5-7). Shott *et al*. systematically evaluated growth patterns in individuals with molecularly-confirmed KS1, revealing postnatal growth retardation in the vast majority (8) and confirming previous reports from clinically diagnosed patients (9-11). Most of the molecularly-confirmed individuals with KS1 did not meet criteria for growth hormone deficiency (12). Despite this, treatment with recombinant growth hormone therapy improved linear growth in some but not all individuals with KS1 (13). Such observations demonstrate the lack of knowledge about the underlying mechanism of growth retardation in KS1 and suggest it is more complex than isolated growth hormone deficiency.

Using morpholinos to knock down *Kmt2d* in zebrafish, Van Laarhoven *et al*. postulated a defect in neural crest-derived cell function in the development of facial flattening in fish deficient in *Kmt2d* because multiple cartilaginous structures were underdeveloped (14). In our previous work we characterized a mouse model of KS1, which exhibits many features seen in patients with the disorder (15). *Kmt2d*^*+/ β Geo*^ mice have disrupted H3K4 trimethylation (H3K4me3) in the dentate gyrus of the hippocampus and associated neurogenesis defects and memory deficits. All three features could be reversed by postnatal administration of agents that favor chromatin opening such as HDAC inhibitors or the HDAC inhibitor-like ketone body, beta-hydroxybutyrate (15, 16). *Kmt2d*^*+/β Geo*^ mice also weigh less than *Kmt2d*^*+/+*^ littermates and exhibit a flattened facial profile like individuals with KS1 (15). The cellular and molecular basis for this phenotype has not been examined.

Here, using high-resolution micro-computed tomography (micro-CT), we elucidate a robust and quantitative skeletal growth retardation phenotype impacting the long bones and cranial base of the skull in *Kmt2d*^*+/ β Geo*^ mice. Histological data from growth plates within both sites suggest a unifying mechanism of disrupted endochondral ossification. Our *in vitro* studies of stable chondrocyte cell lines harboring loss of function *Kmt2d* mutations provide further evidence that increased proliferation and precocious differentiation of chondrocytes play a key role in KS1 pathogenesis. Targeted and genome-wide transcriptome profiling supports precocious differentiation of chondrocytes, and together with targeted chromatin immunoprecipitation studies, suggests the basis for *Sox9* dysregulation. This involves loss of KMT2D-mediated H3K4me3 at an unsuspected target, *Shox2*, causing release of *Sox9* inhibition, and thereby precocious chondrocyte differentiation. Our findings provide novel mechanistic insight into the pathogenesis of growth retardation in KS1 and related disorders and suggest potential novel therapeutic targets.

## Results

### *Kmt2d*^*+/βGeo*^ mice exhibit a specific skeletal growth retardation phenotype

*Kmt2d*^*+/βGeo*^ mice are smaller than *Kmt2d*^*+/+*^ littermates, both grossly (15) and on lateral radiographs (**Figure 1A**). Quantification of body weight (**Figure 1B**) and length (**Figure 1C**) with repeated observations using mice at multiple ages (**Supplemental Figure 1A**) confirmed growth retardation in *Kmt2d*^*+/>β>Geo*^ mice. Because detailed quantification on radiographs revealed a decrease in upper jaw length in *Kmt2d*^*+/β>Geo*^ mice compared to *Kmt2d*^*+/+*^ littermates (data not shown), we first performed high-resolution micro-CT analysis of the skulls of *Kmt2d*^*+/>β>Geo*^ mice and *Kmt2d*^*+/+*^ littermates to examine the craniofacial phenotype. Reconstructed micro-CT images unequivocally confirm a striking flattening of the facial profile in *Kmt2d*^*+/βGeo*^ mice compared to *Kmt2d*^*+/+*^ littermates (**Figure 1D**), which resembles the facial flattening seen in individuals with KS (17).

**Figure 1.**
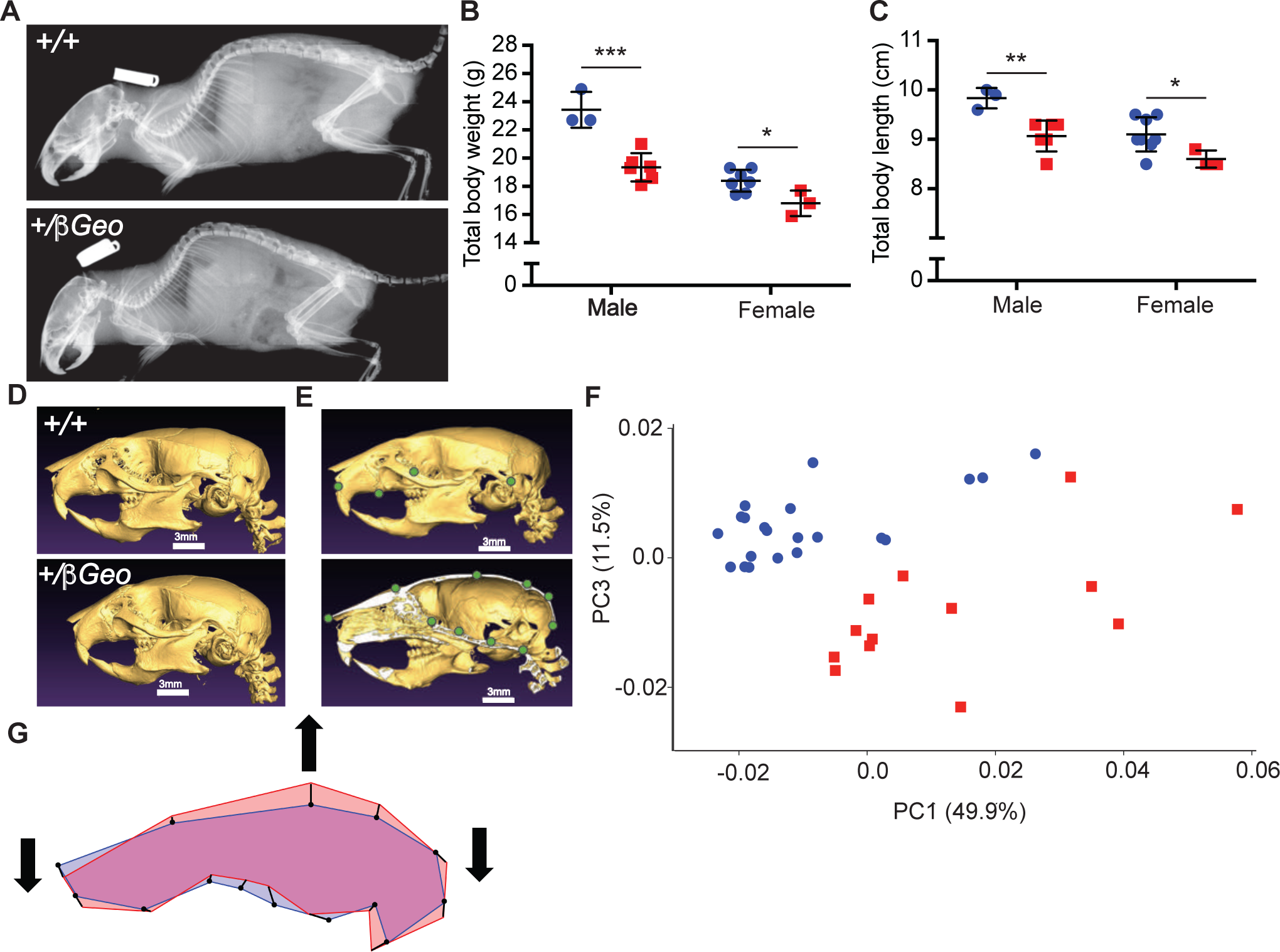
*Kmt2d*^*+/βGeo*^ mice exhibit generalized growth retardation and a specific craniofacial phenotype reminiscent of individuals with KS1. **(A)** Representative radiographs of *Kmt2d*^*+/βGeo*^ mice and *Kmt2d*^*+/+*^ littermates illustrating growth retardation and flattening of the facial profile. Quantification of body weight **(B)** and length **(C)** in 6 week old *Kmt2d*^*+/βGeo*^ male (n=6) and female (n=3) mice and *Kmt2d*^*+/+*^ male (n=3) and female (n=7) littermates. Data represent mean +/-standard deviation, and similar results were obtained with multiple cohorts of mice. Representative reconstructions of high resolution craniofacial micro-CTs in the left lateral view from *Kmt2d*^*+/βGeo*^ mice and *Kmt2d*^*+/+*^ littermates illustrating craniofacial phenotype **(D)** and showing in green 4 pairs of bilateral landmarks (top) and 10 midline landmarks (bottom) used for morphometric analysis **(E)**. **(F)** Principal components analysis of shape revealing separation of two distinct groups along PC1 with *Kmt2d*^*+/+*^ mice (n=21) toward the lower end and *Kmt2d*^*+/βGeo*^ mice (n=13) toward the upper end. **(G)** Overlay of wireframes in left lateral view illustrating relative differences in shape of *Kmt2d*^*+/+*^ mice (blue) and *Kmt2d*^*+/βGeo*^ mice (red). Black vectors show displacement of landmarks associated with the range of shape variation observed on PC1 and indicate ventral bowing, dorsal expansion, and brachycephaly. Thick black arrows illustrate overall shape change in KS1. *Kmt2d*^*+/+*^ mice indicated with blue circles; *Kmt2d*^*+/βGeo*^ mice indicated with red squares. Scale bar=3mm. *p value <0.05; **p value <0.01; ***p value <0.001.

To further characterize the craniofacial phenotype, we performed morphometric analyses of three-dimensional reconstructions of the micro-CT data. Principal components analysis using 18 landmarks **(Figure 1E**) revealed that *Kmt2d*^*+/βGeo*^ and *Kmt2d*^*+/+*^ mice fall into two distinct groups, with *Kmt2d*^*+/β Geo*^ mice toward the upper end of PC1, and approximately half of the total variance across the combined sample being attributable to ventral bowing and brachycephaly (PC1) in *Kmt2d*^*+/βGeo*^ mice (**Figure 1F-G**, **Supplemental Figure 2**). This analysis also indicates that the *Kmt2d*^*+/βGeo*^ mice exhibit a dorsal expansion of the cranial vault compared to *Kmt2d*^*+/+*^ littermates (**Figure 1G and Supplemental Figure 2**).

**Figure 2.**
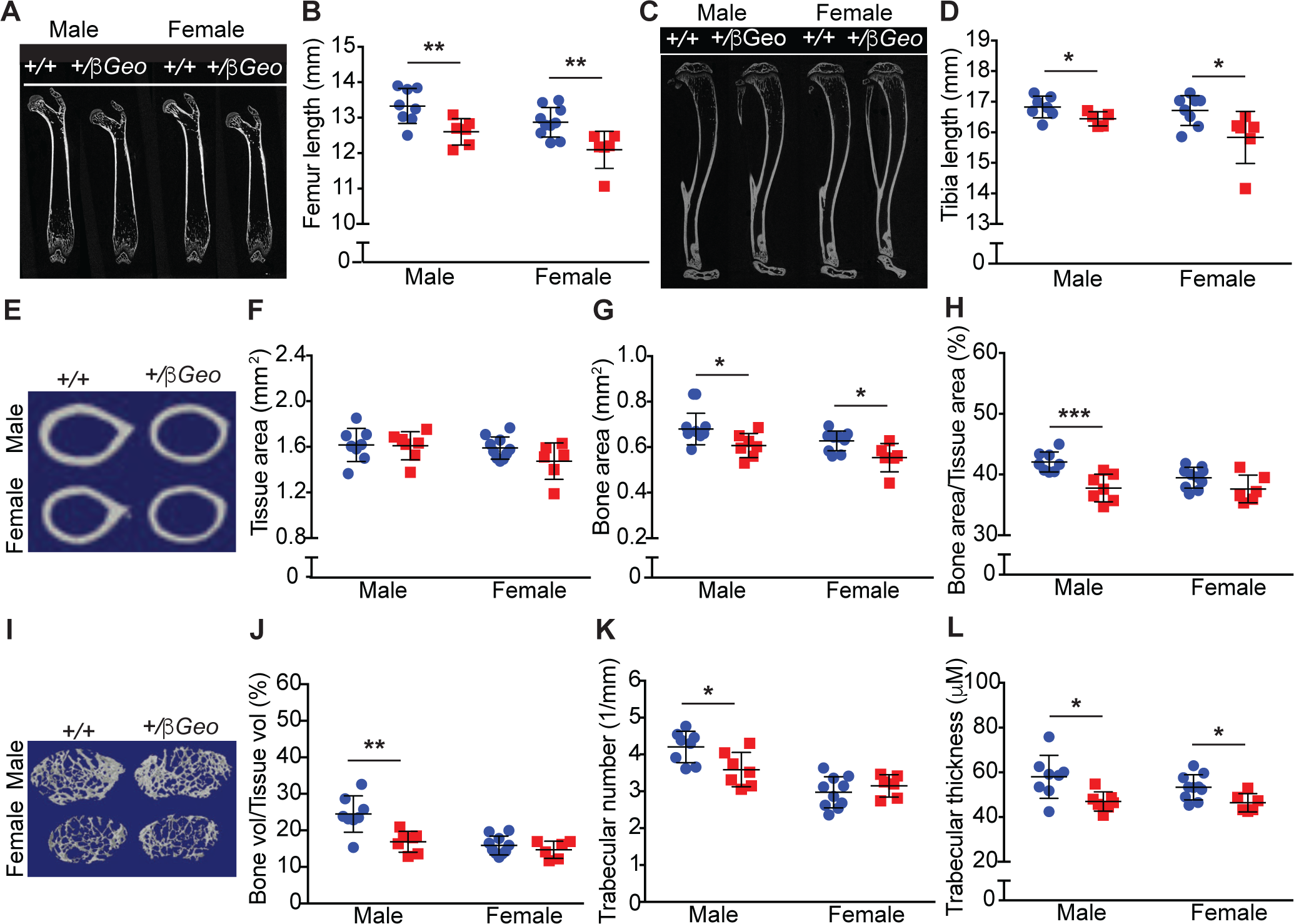
High-resolution micro-CT analysis of long bones in KS1 reveals shortening, thinning, and altered trabecular bone formation in *Kmt2d*^***+/βGeo***^ mice. Femurs **(A, B)** and tibias **(C, D)** are shorter in *Kmt2d*^*+/βGeo*^ mice compared to *Kmt2d*^*+/+*^ littermates. Overall cross-sectional area does not differ between *Kmt2d*^*+/βGeo*^ and *Kmt2d*^*+/+*^ femurs **(E, F)**. Cross-sectional area of mineralized bone is reduced in *Kmt2d*^*+/βGeo*^ femurs compared to *Kmt2d*^*+/+*^ femurs **(E, G)**. The percent of cross-sectional area made up of mineralized bone is reduced in male *Kmt2d*^*+/βGeo*^ femurs compared to *Kmt2d*^*+/+*^ femurs, whereas the difference is not significant in females **(E, H)**. *Kmt2d*^*+/βGeo*^ femurs appear to have decreased trabecular bone near the growth plate compared to *Kmt2d*^*+/+*^ femurs **(I)**. Specifically, percent of tissue volume made up of bone is decreased in male *Kmt2d*^*+/βGeo*^ femurs **(I, J)**, and trabecular number is decreased in male *Kmt2d*^*+/βGeo*^ femurs **(I, K)**. Trabecular thickness is decreased in male and female *Kmt2d*^*+/βGeo*^ femurs **(I, L)**. Blue squares represent *Kmt2d*^*+/+*^ mice (for femurs, n=18; 8 male and 10 female; for tibias n=15; 7 male and 8 female); red circles indicate *Kmt2d*^*+/βGeo*^ mice (for femurs, n=13; 7 male and 6 female, except for femur length where only 6 male mutants could be measured; for tibias, n=11; 5 male, 6 female). Data represent mean +/-standard deviation. Two-sided unpaired student’s t-test was used. *p value <0.05; **p value <0.01; ***p value<0.001.

Because long bones also appeared to be shortened on lateral radiographs (**Figure 1A**) and generalized postnatal growth retardation is a key aspect of the KS skeletal phenotype in patients, we performed analyses of femur and tibial length and high-resolution micro-CT evaluation of bone architecture. Both femurs (**Figure 2A-B**) and tibias (**Figure 2C-D**) were significantly shorter in *Kmt2d*^*+/βGeo*^ mice compared to *Kmt2d*^*+/+*^ littermates at 6 and 18 weeks of age (**Figure 2A-D; Supplemental Figure 1B-C**). High-resolution micro-CT revealed differences in cortical and trabecular bone structure in *Kmt2d*^*+/βGeo*^ mice compared to *Kmt2d*^*+/+*^ littermates. In the femur, cortical cross-sectional tissue area did not differ between groups (**Figure 2E-F**), mineralized bone area was reduced in *Kmt2d*^*+/βGeo*^ mice compared to *Kmt2d*^*+/+*^ littermates (**Figure 2E, G**), which led to a significant decrease in the percent bone area per tissue area in *Kmt2d*^*+/βGeo*^ males (**Figure 2E, H**). Trabecular bone volume examined in the distal femur was reduced in *Kmt2d*^*+/βGeo*^ male mice **(Figure 2I-J)** secondary to decreases in trabecular number (**Figure 2I, K**) and trabecular thickness (**Figure 2I, L**); however in female *Kmt2d*^*+/βGeo*^ mice, only trabecular thickness was significantly reduced **(Figure 2I-L)**. Therefore, *Kmt2d*^*+/βGeo*^ mice exhibit a distinct defect in skeletal growth that is evident at multiple sites and resembles that observed clinically in KS1.

### *In vivo* and *in vitro* studies suggest disrupted endochondral ossification in KS1

Long bones, as well as a few bones in the skull, increase in length by means of endochondral ossification at growth plates, which involves proliferation and hypertrophy of chondrocytes and production of cartilaginous matrix that is ultimately replaced by bone laid down by osteoblasts. The pattern of growth abnormalities in *Kmt2d*^*+/βGeo*^ mice suggests alterations in growth plate dynamics as a potential mechanism for the defects observed in long bones and the cranial base. Therefore we stained longitudinal sections from *Kmt2d*^*+/ βGeo*^ and *Kmt2d*^*+/+*^ proximal tibia growth plates with hematoxylin and eosin (H&E). While overall morphology was intact, we found the growth plates were expanded in *Kmt2d*^*+/βGeo*^ mice compared to *Kmt2d*^*+/+*^ littermates (**Figure 3A**). Growth plate height (**Figure 3A-B**), proliferative zone height (**Figure 3A, C**), and hypertrophic zone height (**Figure 3A, D**) were increased in both male and female *Kmt2d*^*+/βGeo*^ mice compared to *Kmt2d*^*+/+*^ littermates. Counting of the number of chondrocytes per column in the proliferative zone (**Figure 3A, E**) and in the hypertrophic zone (**Figure 3A, F**) revealed increased cell numbers in *Kmt2d*^*+/βGeo*^ mice compared to *Kmt2d*^*+/+*^ mice at both sites, whereas no difference in cell size was evident (**Supplemental Figure 3**). Therefore we examined the cranial base, a key area of the skull that grows by endochondral ossification, and another site which also appears to be shortened in individuals and mice with KS1 (**Figure 1**).

**Figure 3.**
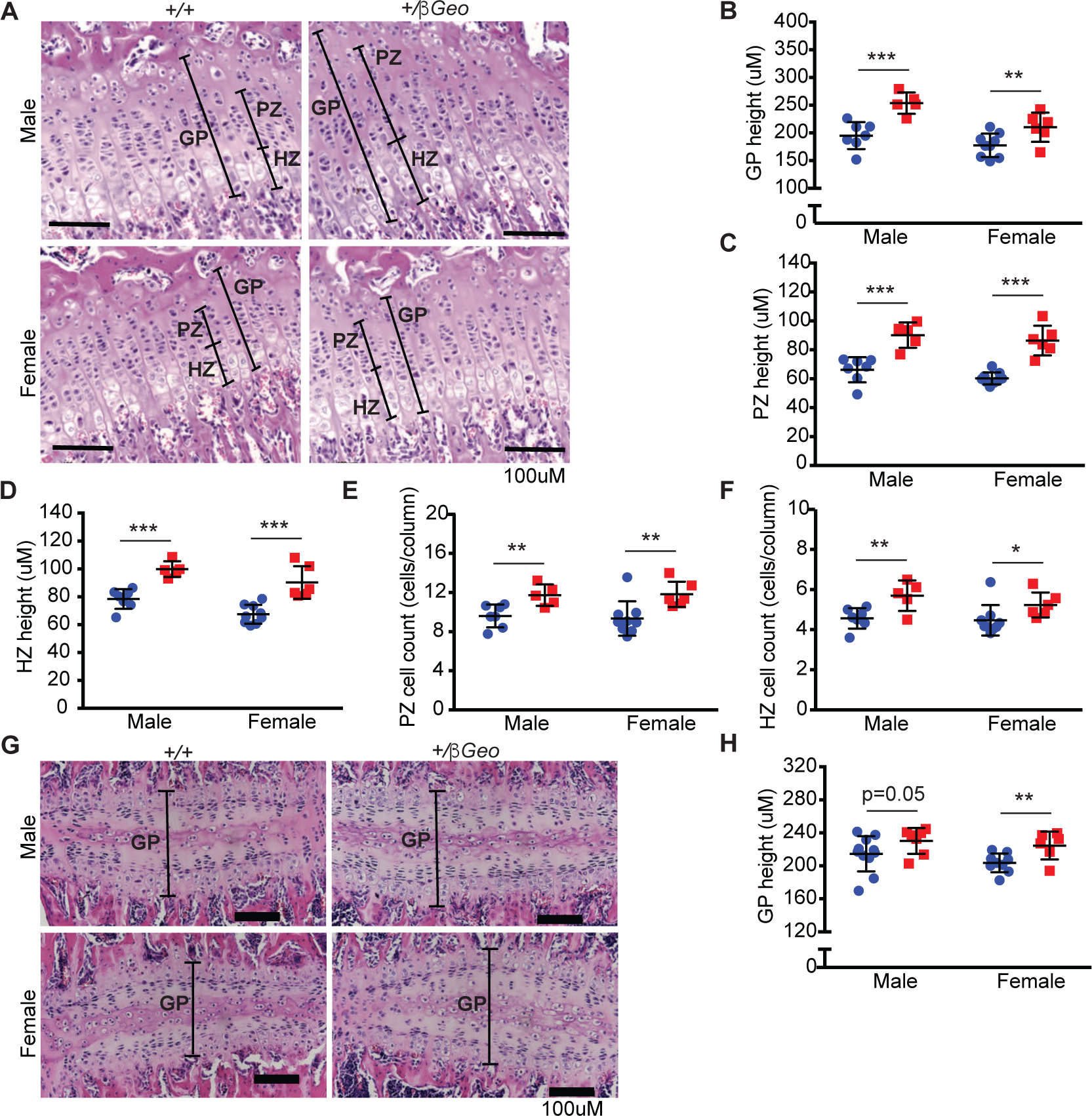
Growth plates from long bones and within the cranial base are expanded in *Kmt2d*^*+/βGeo*^ mice. Proximal tibia growth plates **(A, B)** and their proliferative **(A, C)**, and hypertrophic **(A, D)** zones are expanded in *Kmt2d*^*+/βGeo*^ mice compared to *Kmt2d*^*+/+*^ littermates. The mechanism involves increased cell numbers per column in both the proliferative **(E)** and hypertrophic **(F)** zones. **(G, H)** Cranial base intersphenoidal synchondroses from *Kmt2d*^*+/βGeo*^ mice are expanded compared to *Kmt2d*^*+/+*^ littermates. GP, growth plate; PZ, proliferative zone; HZ, hypertrophic zone. Blue circles represent *Kmt2d*^*+/+*^ mice (n=16 proximal tibia growth plates; 7 male and 9 female; n=20 intersphenoidal synchondroses (ISS); 11 male, 9 female); red squares represent *Kmt2d*^*+/βGeo*^ mice (n=11 proximal tibia growth plates; 5 male and 6 female; n=13 ISS’s; 7 male and 6 female). Data represent mean +/-standard deviation. *p value <0.05; **p value <0.01; ***p value<0.001.

Examination of intrasphenoidal synchondroses, which are partly responsible for growth of the cranial base in the anterior-posterior direction (18), revealed a similar expansion in *Kmt2d*^*+/ βGeo*^ mice compared to *Kmt2d*^*+/+*^ littermates (**Figure 3G-H**). While counterintuitive that growth plate height could be expanded whilst the cranial base and long bones themselves were shortened in length, these findings suggest that abnormal endochondral ossification of long bones and of the cranial base are a unifying mechanism for the skeletal phenotype of KS1 mice.

To understand the cellular basis for this phenotype, we employed a more manipulatable system of chondrocyte development, the well-established ATDC5 cell system; upon induction, these mesenchymal progenitors progress through the stages of chondrocyte development, including proliferation and hypertrophy (19). Using CRISPR-Cas9 genome editing technology, we created stable cell lines with homozygous mutations of varying severity, *Kmt2d*^*ΔR5551/-*^, which has a deletion of a single amino acid on one allele and deletion of the SET domain on the other allele, and *Kmt2d*^*-/-*^, which has bi-allelic deletions of the SET domain, and compared them to vector only parental ATDC5 cells (*Kmt2d*^*+/+*^ ; **Figure 4; Supplemental Figure 4**) before and after induction of chondrocyte differentiation. Assessment of cellular proliferation by the MTT assay (**Supplemental Figure 5A**), and direct cell counting (**Supplemental Figure 5B**) indicated that loss of *Kmt2d* function increases chondrocyte replication in accordance with the severity of the mutation. This was evident at 4-7 days post-differentiation induction. These findings support our previous observation of proliferative zone expansion in proximal tibia growth plates from *Kmt2d*^*+/βGeo*^ mice (**Figure 3A, C, E**). Similarly, cultures of *Kmt2d*^*-/-*^ cells exhibited more dramatic increases in matrix deposition, as indicated by alcian blue staining, when compared to *Kmt2d*^*+/+*^ controls with *Kmt2d*^*ΔR5551/-*^ cells having an intermediate phenotype (**Figure 4A-B**). By fourteen days after induction of differentiation, we observed a significant increase in alcian blue staining of *Kmt2d*^*-/-*^ chondrocytes compared to *Kmt2d*^*Δ R5551/-*^ and *Kmt2d*^*+/+*^ chondrocytes (**Figure 4A-B**). This increase was progressive and exceedingly apparent at day 21. These data suggest precocious differentiation of *Kmt2d*^*-/-*^ cells and fits with the expansion of hypertrophic zones observed in proximal tibia growth plates from *Kmt2d*^*+/βGeo*^ mice compared to *Kmt2d*^*+/+*^ littermates (**Figure 3A, D, F**). Together, these findings show that all cell lines (*Kmt2d*^*+/+*^, *Kmt2d*^*ΔR5551/-*^ and *Kmt2d*^*-/-*^) proceed through the phases of chondrocyte development, i.e proliferation (days 4-7; **Supplemental Figure 5A-B**) and hypertrophy (days 14-21; **Figure 4A-B**); however, *Kmt2d*^*-/-*^ chondrocytes exhibit increased proliferation and precocious differentiation compared to *Kmt2d*^*+/+*^ (and in some cases *Kmt2d*^*ΔR5551/-*^) cells (**Figure 4A-B; Supplemental Figure 5A-B**). These observations support our *in vivo* findings of growth plate expansion in both tibias and cranial base intrasphenoidal synchondroses of *Kmt2d*^*+/βGeo*^ mice compared to *Kmt2d*^*+/+*^ littermates (**Figure 3**), and together these data support increased proliferation and precocious differentiation as unifying features of KMT2D-deficient chondrocytes both *in vitro* and *in vivo.*

**Figure 4.**
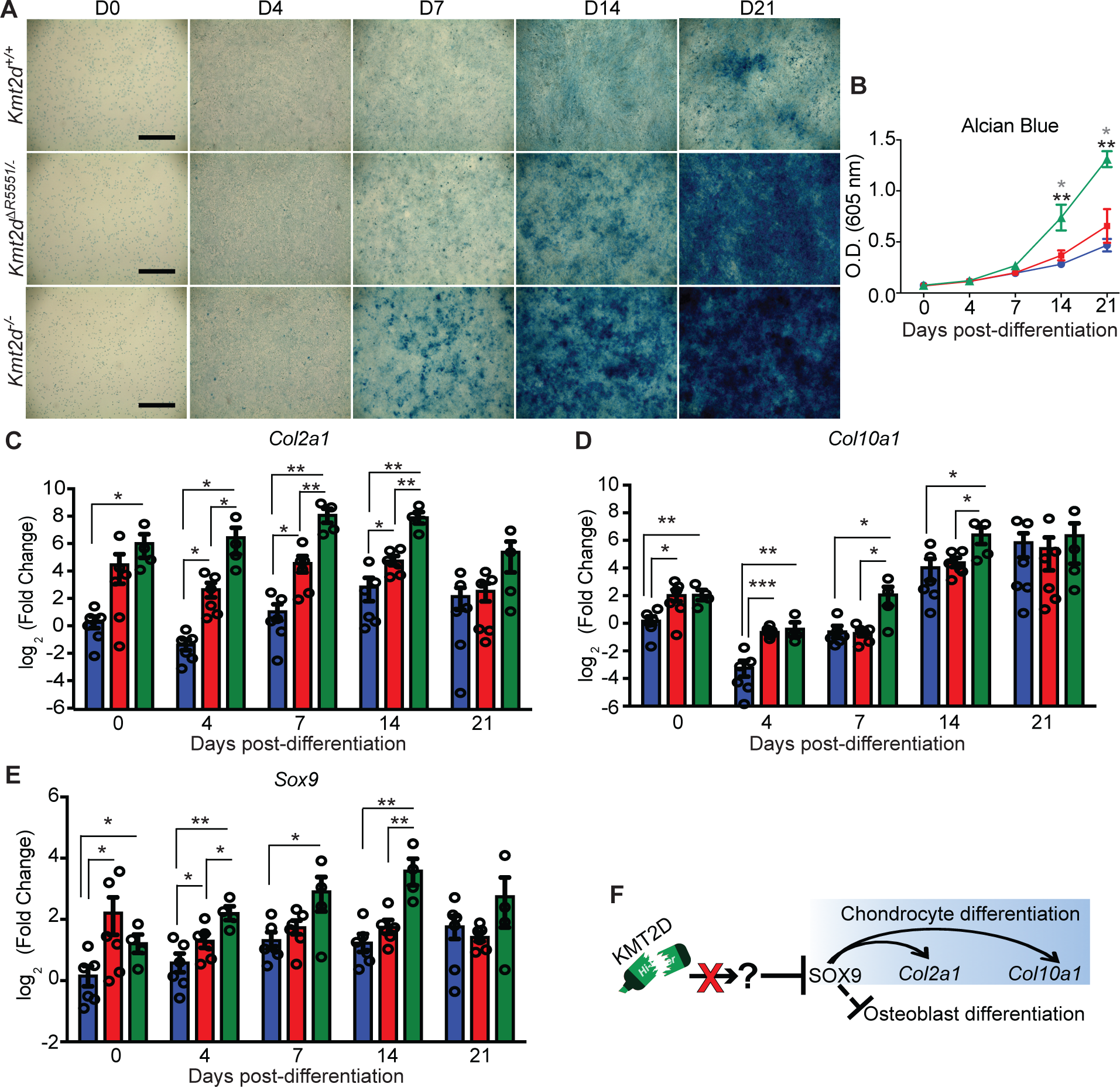
Precocious differentiation of *Kmt2d*^*-/-*^ and *Kmt2d*^*ΔR5551/-*^ chondrocytes. Stable ATDC5 cell lines with *Kmt2d* mutations of varying severity were created using CRISPR/Cas9 genome editing technology. Chondrocyte differentiation was induced at day 0. Alcian blue staining was used to visualize **(A)** and quantify **(B)** chondrocyte differentiation over time. Quantitative RT-PCR with primers specific for *Col2a1* **(C)**, *Col10a1* **(D)**, and *Sox9* **(E)** was performed on RNA isolated from undifferentiated (day 0) and differentiated *Kmt2d*^*+/+*^, *Kmt2d*^*Δ R5551/-*^, and *Kmt2d*^*-/-*^ stable chondrocyte cell lines at 4, 7, 14, and 21 days after induction of differentiation. Fold change was calculated relative to undifferentiated *Kmt2d*^*+/+*^ cells (day 0). Blue bars represent *Kmt2d*^*+/+*^ cells, red bars represent *Kmt2d*^*Δ R5551/-*^ cells, and green bars represent *Kmt2d*^*-/-*^ cells. Error bars indicate standard error of the mean**. (F)** Preliminary model for precocious chondrocyte differentiation in KS1. Scale bar =1mM. Black asterisks represent differences between *Kmt2d*^*+/+*^ and *Kmt2d*^*-/-*^ cells while the gray asterisks represent differences between *Kmt2d*^*-/-*^ and *Kmt2d*^*Δ R5551/-*^. *p value <0.05; **p value <0.01; ***p value<0.001.

**Figure 5.**
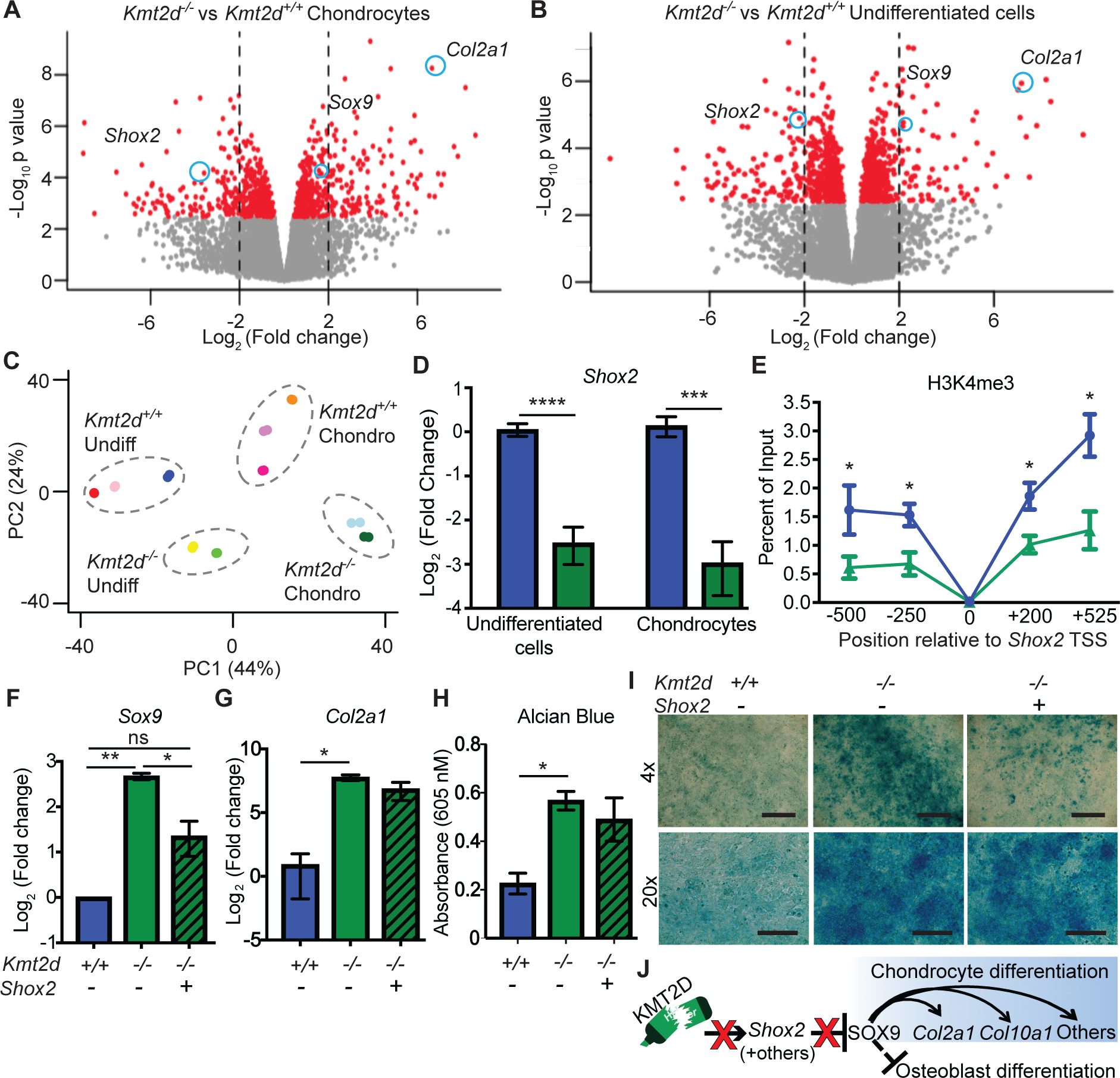
Genome-wide transcriptome profiling identifies *Shox2* as a novel target of KMT2D-mediated H3K4me3 and mediator of precocious chondrocyte differentiation in KS1. Volcano plots showing genome-wide differential gene expression in *Kmt2d*^*-/-*^ and *Kmt2d*^*+/+*^ chondrocytes **(A)** and undifferentiated cells **(B).** Principal components analysis **(C)** reveals tight clustering within and distinct separation between genotypes and differentiation states, separating the cells into 4 distinct groups. *Kmt2d*^*+/+*^ cells cluster toward the upper end of PC2 and *Kmt2d*^*-/-*^ cells cluster toward the lower end of PC2. Undifferentiated cells cluster toward the lower end of PC1 and chondrocytes cluster toward the upper end of PC1. Each color represents a biological replicate; each biological replicate is a distinct clonal cell line. Three (*Kmt2d*^*+/+*^) or two (*Kmt2d*^*-/-*^) biological replicates were used for each differentiation state (chondrocytes vs undifferentiated cells). Each point represents a technical replicate; two technical replicates were performed for each cell line. *Sox9 and Col2a1* are upregulated in *Kmt2d*^*-/-*^ chondrocytes (**A**) and *Kmt2d*^*-/-*^ undifferentiated cells **(B)**, and *Shox2* is down-regulated in *Kmt2d*^*-/-*^ chondrocytes (**A**) and *Kmt2d*^*-/-*^ undifferentiated cells **(B).** *Shox2* down-regulation was validated with qPCR in three independent experiments **(D)** with blue bars representing *Kmt2d*^*+/+*^ cells and green bars representing *Kmt2d*^*-/-*^ cells. **(E)** Chromatin Immunoprecipitation followed by qPCR (ChIP-qPCR) revealed decreased H3K4me3 along the *Shox2* promoter in *Kmt2d*^*-/-*^ cells (green) compared to *Kmt2d*^*+/+*^ cells (blue). Overexpression of *Shox2* in *Kmt2d*^*-/-*^ cells led to partial recovery (downregulation) of *Sox9* expression **(F)** but did not lead to significant recovery of *Col2a1* expression **(G)** or alcian blue staining **(H-I)**. **(J)** Model for molecular pathogenesis of KS1 growth retardation. Error bars represent standard error of the mean. Scale bars=1mm (top panels) and 200μM (bottom panels). *p value <0.05; **p value <0.01; ***p value<0.001; ****p<0.0001. Chondro, chondrocytes; Undiff, undifferentiated cells.

### Precocious chondrocyte expression patterns in a cellular model of KS1

Using qRT-PCR, we measured expression of chondrocyte differentiation markers *Col2a1* and *Col10a1* in undifferentiated mesenchymal cells and in differentiated chondrocytes (**Figure 4C-D**). *Col2a1* is normally expressed early in the chondrocyte developmental program, initially within the proliferative phase, and *Col10a1* is expressed later in more advanced stages of chondrocyte development beginning in the early hypertrophic phase (20). Expression of *Col2a1* was increased in undifferentiated *Kmt2d*^*-/-*^ cells compared to undifferentiated *Kmt2d*^*+/+*^ cells (**Figure 4C**; day 0). Upon differentiation induction, we observed an initial decrease in *Col2a1* expression in *Kmt2d*^*+/+*^ and *Kmt2d*^*ΔR5551/-*^ cells, followed by a striking and progressive dose-dependent increase in *Col2a1* expression in *Kmt2d*^*ΔR5551/-*^ and *Kmt2d*^*-/-*^ chondrocytes compared to *Kmt2d*^*+/+*^ chondrocytes (**Figure 4C**). While a gradual increase in *Col2a1* expression is expected and was observed over time (days 4-14) in *Kmt2d*^*+/+*^ chondrocytes, in *Kmt2d*^*ΔR5551/-*^ and *Kmt2d*^*-/-*^ chondrocytes, *Col2a1* expression increased precociously, attaining higher maximal levels earlier – by day 7 compared to day 14 in *Kmt2d*^*+/+*^ cells (**Figure 4C**). Similar to *Col2a1*, expression of the later *Col10a1* chondrocyte differentiation marker was increased in undifferentiated *Kmt2d*^*Δ R5551/-*^ and *Kmt2d*^*-/-*^ cells compared to undifferentiated *Kmt2d*^*+/+*^ cells (day 0; **Figure 4D**). Upon induction of chondrocyte differentiation in *Kmt2d*^*+/+*^ cells, expression of *Col10a1* initially dropped to almost undetectable levels; however by day 14, we observed a 16-fold increase in *Col10a1* expression above baseline levels with maximal expression at day 21 (**Figure 4D**), consistent with *Col10a1* being a late marker of hypertrophic chondrocyte differentiation (21). Over time both *Kmt2d*^*ΔR5551/-*^ and *Kmt2d*^*-/-*^ chondrocyte cell lines showed a pattern similar to *Kmt2d*^*+/+*^ with an initial decrease in *Col10a1* expression, followed by a robust increase in expression (**Figure 4D**). However *Col10a1* expression remained 4-fold higher in *Kmt2d*^*-/-*^ cells compared to *Kmt2d*^*+/+*^ cells through day 14, and the maximal level of expression was attained precociously at day 14 in *Kmt2d*^*-/-*^ chondrocytes compared to day 21 in *Kmt2d*^*+/+*^ chondrocytes (**Figure 4D**). *Kmt2d*^*ΔR5551/-*^ cells showed intermediate levels of expression overall, resembling *Kmt2d*^*-/-*^ cells until day 4 and *Kmt2d*^*+/+*^ cells thereafter (**Figure 4D**).

### Overexpression of *Sox9* mediates the precocious differentiation phenotype

We hypothesized that *Sox9*, which encodes a key transcription factor that promotes chondrocyte differentiation and is a known direct activator of *Col2a1* and *Col10a1* (20), could be a potential candidate gene to mediate our cellular phenotype. SOX9 facilitates chondrocyte differentiation through proliferation and hypertrophy and prevents further progression toward bone-forming osteoblasts (20). The level of expression of *Sox9* was low in undifferentiated *Kmt2d*^*+/+*^ cells; after chondrocyte differentiation was induced, *Sox9* expression gradually increased by 3-4-fold over 21 days (**Figure 4E**). Both *Kmt2d*^*ΔR5551/-*^ and *Kmt2d*^*-/-*^ cells showed increased *Sox9* expression relative to *Kmt2d*^*+/+*^ cells in the undifferentiated state (**Figure 4E**). Upon differentiation, we observed an increase in *Sox9* expression dependent upon the severity of the KMT2D mutation (**Figure 4E**). Like *Col2a1* and *Col10a1*, which are induced by SOX9, *Sox9* is expressed at significantly higher levels in *Kmt2d*^*-/-*^ compared to *Kmt2d*^*+/+*^ chondrocytes up to 14 days after differentiation induction. For *Sox9*, this culminates in a striking 4-fold increase in expression in *Kmt2d*^*-/-*^ chondrocytes compared to *Kmt2d*^*+/+*^ chondrocytes at day 14 (**Figure 4E**). The observation that genetic ablation of KMT2D, a histone methyltransferase writer of an activating mark, increases *Sox9* suggests a model by which loss of KMT2D may prevent expression of an inhibitor of *Sox9*, thus allowing subsequent chondrocyte differentiation to proceed precociously and at the expense of osteoblast differentiation (**Figure 4F**).

### Loss of *Shox2* expression and release of *Sox9* inhibition in a KS1 chondrocyte model

We performed genome-wide transcriptome profiling by RNA-seq using RNA from *Kmt2d*^*-/-*^ and *Kmt2d*^*+/+*^ chondrocytes 7 days after induction of differentiation to identify KMT2D targets that can inhibit *Sox9* (**Figure 5A; Supplemental Figure 6A; Supplemental Table S1**). Day 7 appears to be a key time point for gene expression changes that lead to precocious differentiation because immediately after day 7 alcian blue staining increases significantly in *Kmt2d*^*-/-*^ chondrocytes compared to *Kmt2d*^*+/+*^ chondrocytes (**Figure 4A**,**B**), and just prior to day 7 we observed a striking difference in expression of *Sox9* between *Kmt2d*^*-/-*^ and *Kmt2d*^*+/+*^ chondrocytes (**Figure 4E**). For comparison, we also performed RNA-seq on undifferentiated *Kmt2d*^*-/-*^ and *Kmt2d*^*+/+*^ mesenchymal cells (**Figure 5B, Supplemental Figure 6B; Supplemental Table S2**).

Multiple observations emerged from our transcriptome profiling. First, principal components analysis revealed highly correlative data with tight clustering within and distinct separation between genotypes and differentiation states, separating the cells into 4 distinct groups (*Kmt2d*^*+/+*^ undifferentiated cells, *Kmt2d*^*+/+*^ chondrocytes, *Kmt2d*^*-/-*^ undifferentiated cells, and *Kmt2d*^*-/-*^ chondrocytes) with differentiation state (chondrocytes versus undifferentiated cells) explaining a greater percentage of the variance than genotype (*Kmt2d*^*-/-*^ versus *Kmt2d*^*+/+*^; **Figure 5C**). Second, differential expression analysis revealed multiple expected gene expression changes, including greater than 50-fold upregulation of *Col2a1* in *Kmt2d*^*-/-*^ cells compared to *Kmt2d*^*+/+*^ cells, both in chondrocytes (day 7; **Figure 5A; Supplemental Figure 6A; Supplemental Table S1**) and in the undifferentiated state (day 0; **Figure 5B**; **Supplemental Figure 6B; Supplemental Table S2; Tables 1-2**), similar to our previous observations (**Figure 4C**). In addition, we observed consistent 3-4 fold upregulation of the master transcriptional regulator of chondrogenesis, *Sox9*, in *Kmt2d*^*-/-*^ cells compared to *Kmt2d*^*+/+*^ cells in both differentiation states (**Figure 5A-B; Supplemental Tables S1-3**), again supporting our previous findings (**Figure 4E**). Additional well-established markers of chondrocyte differentiation(22) were upregulated as well in *Kmt2d*^*-/-*^ cells compared to *Kmt2d*^*+/+*^ cells, including *Col11a1* (22; over 10-fold in both chondrocytes and undifferentiated cells; **Supplemental Tables S1-S4**), *Col9a1* (22; over 10-fold in chondrocytes; **Supplemental Table S1**), and *Col10a1* (4-fold in undifferentiated cells; **Supplemental Table S2;** compare to **Figure 4D**), further supporting a mechanism involving precocious chondrocyte differentiation. Lastly, we observed more differential gene expression within a single genotype, *Kmt2d*^*+/+*^ (7,201 genes; **Supplemental Table S5**) or *Kmt2d*^*-/-*^ (4,764 genes; **Supplemental Table S6**), over the course of differentiation from mesenchymal cells to chondrocytes compared to the differential expression observed between genotypes within a particular differentiated state, i.e. chondrocytes *Kmt2d*^*-/-*^ versus *Kmt2d*^*+/+*^ (932 genes; **Supplemental Table S1)** or undifferentiated mesenchymal cells *Kmt2d*^*-/-*^ versus *Kmt2d*^*+/+*^ (1009 genes; **Supplemental Table S2)**. This is expected and suggests that the number of gene expression changes required for a cell to undergo a differentiation program is greater than the number that result from loss of function of a single gene, even if that gene is a component of the epigenetic machinery with many downstream direct targets and indirect consequences.

Comparing *Kmt2d*^*-/-*^ chondrocytes to *Kmt2d*^*+/+*^ chondrocytes, we found 932 differentially expressed genes with 486 upregulated and 446 downregulated (**Figure 5A; Supplemental Figure 6A; Supplemental Table S1**). There were 1009 differentially expressed genes when *Kmt2d*^*-/-*^ and *Kmt2d*^*+/+*^ undifferentiated cells were compared with 474 being upregulated and 535 being downregulated (**Figure 5B; Supplemental Figure 6B; Supplemental Table S2**). Of those differentially expressed genes, 382 are common across differentiation state (i.e. found in both chondrocytes and undifferentiated cells; **Supplemental Table S3**), and this represents a highly significant overrepresentation (Fisher’s test, p < 2.2e10 ^-16^, odds ratio = 14.2). Of the 932 differentially expressed genes in chondrocytes, 265 were differentially expressed at least 4-fold (**Figure 5A, Supplemental Figure 6A**), and of the 1009 differentially expressed genes in undifferentiated cells, 180 were differentially expressed least 4-fold (**Figure 5B, Supplemental Figure 6B**). 78 are common across chondrocytes and undifferentiated cells **(Supplemental Table S4)**, and this overlap is again highly significant (Fisher’s test, p < 2.2e10 ^-16^, odds ratio = 78.1). Of those 78 differentially expressed transcripts (≥4-fold) common to both chondrocytes and undifferentiated cells, 74 had established gene names. Of those 74 genes, 30 were upregulated and 44 were downregulated in *Kmt2d*^*-/-*^ cells **(Supplemental Table S4)**. We hypothesized that direct KMT2D targets would have more dramatic gene expression abnormalities and thus focused on the subset that were downregulated at least 4-fold in *Kmt2d*^*-/-*^ cells **(Supplemental Table S4)**. We searched the literature to identify whether any of the 44 genes had previously been reported as targets of KMT2D, regulators (particularly inhibitors) of *Sox9*, and/or involved in chondrocyte development. No known KMT2D targets were among the 44 genes identified by our transcriptome analysis to be down-regulated at least 4-fold in *Kmt2d*^*-/-*^ cells. Three genes were identified as being involved in chondrocyte development and connected with *Sox9* in the literature; however, only one had the potential to negatively regulate *Sox9* based on previous studies *–* the transcription factor *Shox2.*

*Shox2* is the most intriguing candidate gene down-regulated over 4-fold **(Supplemental Table S4)** in our genome-wide transcriptome analysis comparing *Kmt2d*^*-/-*^ cells to *Kmt2d*^*+/+*^ cells in differentiated chondrocytes (**Figure 5A; Supplemental Figure 6A**) and undifferentiated cells (**Figure 5B; Supplemental Figure 6B**). *Shox2* is the mouse paralogue to the human *SHOX* gene, which when disrupted causes short stature (23-25). *Shox2* is important in mice for long bone growth and chondrocyte development (26,27), and importantly, conditional deletion in chondrocytes leads to precocious differentiation with concomitant increase in *Col2a1* and *Sox9* expression (26), similar to what we observed in *Kmt2d*^*-/-*^ cells (**Figure 4C,E**). We confirmed that *Shox2* is downregulated over 4-fold in *Kmt2d*^*-/-*^ cells compared to *Kmt2d*^*+/+*^ cells in the undifferentiated state and in chondrocytes by qPCR in three independent experiments (**Figure 5D**).

### Functional testing supports the idea that *Shox2* plays a mechanistic role in the cellular differentiation phenotype

Based on these observations, we hypothesized *Shox2* may be a target of KMT2D and a mediator of the precocious differentiation observed in *Kmt2d*^*-/-*^ cells. KMT2D places the H3K4me3 activating mark at gene promoters (28), and we previously showed depletion of this mark in the hippocampi of *Kmt2d*^*+/ βGeo*^ mice (15). Moreover, examination of ENCODE data from mouse limb bud revealed a strong peak of H3K4me3 at the 5’ end of the *Shox2* gene (29, 30); **Supplemental Figure 7A**). We therefore performed chromatin immunoprecipitation followed by quantitative PCR (ChIP-qPCR) using an antibody that specifically recognizes H3K4me3 in *Kmt2d*^*-/-*^ and *Kmt2d*^*+/+*^ cells. We observed a significant depletion of H3K4me3 at multiple sites within this region in *Kmt2d*^*-/-*^ cells compared to *Kmt2d*^*+/+*^ cells (**Figure 5E; Supplemental Figure 7B-F**). In contrast, H3 levels were no different between any of the cell lines **(Supplemental Figure 7B-F**). This suggests that when present, KMT2D may bind to the *Shox2* promoter and place the H3K4me3 activating mark in these cells. However when KMT2D is absent, less of this mark is placed. Reduction of this mark (as opposed to complete loss) makes sense given the redundancy of the H3K4 histone methyltransferase system. Specifically, reduction of the mark at the *Shox2* promoter in *Kmt2d*^*-/-*^ cells along with decreased *Shox2* expression implicates KMT2D and H3K4me3 in the activation of *Shox2* transcription, providing mechanistic insight into precocious chondrocyte differentiation and the epigenetic regulation thereof.

Next, we wanted to determine whether overexpression of *Shox2* in *Kmt2d*^*-/-*^ cells was sufficient to inhibit precocious chondrocyte differentiation. Transduction of *Kmt2d*^*+/+*^ and *Kmt2d*^*-/-*^ cells with lentivirus overexpressing *Myc*-tagged *Shox2* led to increased *Shox2* transcript levels and increased Myc-tagged SHOX2 protein in one *Kmt2d*^*+/+*^ and one *Kmt2d*^*-/-*^ cell line **(Supplemental Figure 8A-B**). We first examined *Sox9* transcript levels to determine whether overexpression of *Shox2* could restore proper *Sox9* expression in *Kmt2d*^*-/-*^ cells. Similar to previous observations in non-transduced cells, *Kmt2d*^*-/-*^ cells transduced with control virus showed higher *Sox9* transcript levels compared to *Kmt2d*^*+/+*^ cells transduced with control virus. Notably, upon transduction of *Kmt2d*^*-/-*^ cells with *Myc*-tagged *Shox2*, we observed reduced *Sox9* transcript levels, which were not significantly different than those observed in *Kmt2d*^*+/+*^ cells (**Figure 5F**). These results indicate that overexpression of *Shox2* is sufficient to inhibit and therefore restore *Sox9* expression to near wild type levels and supports a key role for SOX9 in precocious chondrocyte differentiation in KS1.

To determine if overexpression of *Shox2* can fully restore the cellular differentiation phenotype in *Kmt2d*^*-/-*^ cells, we looked at *Col2a1* expression and alcian blue staining. Similar to previous observations in non-transduced cells, *Kmt2d*^*-/-*^ cells transduced with control virus showed higher *Col2a1* transcript levels and alcian blue staining compared to *Kmt2d*^*+/+*^ cells transduced with control virus. Upon transduction of *Kmt2d*^*-/-*^ cells with *Myc*-tagged *Shox2*, we did not observe a significant difference in *Col2a1* expression or alcian blue staining between *Kmt2d*^*-/-*^ cells overexpressing *Shox2* and *Kmt2d*^*-/-*^ cells transduced with control virus, nor did we observe a significant difference in these parameters when we compared *Kmt2d*^*-/-*^ cells overexpressing *Shox2 and Kmt2d*^*+/+*^ cells transduced with control virus (**Figure 5G-I)**. Rather, we observed intermediate *Col2a1* expression and alcian blue staining upon overexpression of *Shox2* in *Kmt2d*^*-/-*^ cells (**Figure 5G-I)**. These findings suggest an intermediate differentiation state that is not significantly different from *Kmt2d*^*+/+*^ or *Kmt2d*^*-/-*^ cells and moreover that additional cellular targets of KMT2D and of SOX9 contribute to the precocious chondrocyte differentiation observed in KS1 cells. Based on these data, our model for precocious chondrocyte differentiation-mediated growth retardation in KS is that deficiency of *Kmt2d*-mediated H3K4me3 leads to impaired activation of *Shox2*, among other targets **(Figure 5J)**. This allows for *Sox9* overexpression and subsequent precocious chondrocyte differentiation via activation of *Col2a1, Col10a1*, and other chondrocyte-specific genes, and this may occur at the expense of osteoblast differentiation **(Figure 5J)**, leading to the observed skeletal phenotype associated with Kabuki syndrome.

## Discussion

We show for the first time that *Kmt2d*^*+/βGeo*^ mice exhibit a highly specific and quantitative skeletal growth retardation phenotype consisting of decreased body length and weight; shortened long bones; and ventral bowing, dorsal expansion, and brachycephaly of skulls. Our findings resemble those observed classically in individuals with KS, including post-natal short stature and flattened facial profile, and are further supported by more recent detailed observations that patients with *KMT2D* mutations have relatively increased cranial heights and disproportionately shortened tibias (31). This suggests that our *Kmt2d*^*+/βGeo*^ mice are a good model of the human KS1 skeletal growth retardation phenotype and that our findings provide relevant outcome measures for preclinical therapeutic trials in mice and future clinical studies in humans.

This pattern of skeletal features can be explained at least in part by a common mechanism of disrupted endochondral ossification at distinct sites in *Kmt2d*^*+/βGeo*^ mice, namely at long bone growth plates and intrasphenoidal synchondroses within the cranial base. Others have observed similar and additional craniofacial features in a mouse model of the closely related disorder Kabuki syndrome 2 (KS2) due to constitutive and neural crest-specific ablation of the H3K27 demethylase *Kdm6a/Utx* (32). Our findings and theirs could indicate that KS1 and KS2 represent distinct disorders with distinct mechanisms at play. However we favor the idea that common mechanisms may be present in KS1 and KS2, and the seemingly disparate findings are simply due to timing and experimental design. Shpargel *et al.* examined the explicitly neural-crest derived viscerocranium/facial skeleton during embryonic development (32), while our studies focus on the ongoing development of the cranial base in the adult animal. However, both likely influence one another, and the impacts of each will be difficult to parse out. Perhaps, simultaneous ongoing decreased cartilaginous growth of the cranial base as suggested here in KS1, which leads to ventral bowing, dorsal expansion, and brachycephaly of the skull, might contribute to or exacerbate a neural crest-derived craniofacial phenotype previously observed in KS2 (32), but also likely present in KS1. Additional studies will be required to fully understand the pathogenesis of the craniofacial phenotype in KS1 and KS2.

Here we implicate SOX9 in the pathogenesis of the KS1 skeletal phenotype. Our initially unexpected observations of growth plate expansion in the setting of decreased growth at distinct sites in KS1 mice suggested disrupted endochondral ossification as a mechanism and the chondrocyte as a relevant cell type. A more dynamic system of chondrocyte development, stable ATDC5 cell lines with *Kmt2d* loss of function mutations, confirmed precocious chondrocyte differentiation at the cellular and gene expression levels, thereby suggesting a cell-autonomous phenotype reconstituted in an isogenic model. Our findings point to a novel mechanism of loss of KMT2D-mediated release of inhibition of *Sox9*, leading to unchecked and precocious chondrocyte differentiation. Supporting our findings, *Sox9* is a well-known mediator of chondrocyte differentiation required for cartilage formation (33-35). Monoallelic loss of function mutations in humans lead to campomelic dysplasia, a severe chondrodysplasia (36, 37), and heterozygous ablation in mice resembles the human phenotype (34). Conditional ablation in differentiated chondrocytes disrupts endochondral ossification resulting in severely shortened growth plate heights and reveals that *Sox9* expression is necessary for chondrocyte proliferation and late stages of hypertrophy (20). This is consistent with our findings of both increased chondrocyte proliferation and precocious differentiation (hypertrophy) in *Kmt2d*^*-/-*^ stable cell lines in the setting of increased expression of *Sox9.* SOX9 is known to directly activate multiple chondrocyte*-*specific factors, including early and late markers of chondrogenesis, like *Col2a1* and *Col10a1*, respectively (20, 33), which are also dysregulated and may be a measure of SOX9’s down-stream effects although it is unlikely that these are the only genes dysregulated secondary to increased *Sox9* expression. Moreover, loss of KMT2D leading to release of *Sox9* inhibition and subsequent activation of early and late effectors of chondrogenic differentiation would explain our observed increases in proliferative zone, hypertrophic zone, and overall growth plate heights in *Kmt2d*^*+/βGeo*^ mice.

In addition to facilitating chondrocyte differentiation, *SOX9* also prevents osteoblast differentiation (20). The net effect of this imbalance between precocious chondrocyte differentiation and reduced osteoblast differentiation may lead to growth retardation in KS1, specifically to shortening of the long bones and decreased bone formation in *Kmt2d*^*+/βGeo*^ mice. In support of this, when we measured osteoblast differentiation by alizarin red staining, we saw that newly formed bone was decreased in *Kmt2d*^*+/βGeo*^ mouse bone marrow mesenchymal stem cells compared to cells derived from *Kmt2d*^*+/+*^ mice (data not shown). However, future conditional studies will be required to reveal the exact impact in individual lineages.

We used genome-wide transcriptome profiling by RNA-seq to identify KMT2D target genes and associated molecular pathways that may play a role in KS1 pathogenesis. We identified *Shox2* as a novel target of KMT2D. *Shox2* was downregulated over 4-fold in *Kmt2d*^*-/-*^ chondrocytes and undifferentiated mesenchymal cells. Depletion of the H3K4me3 activating mark placed by KMT2D at the *Shox2* promoter in *Kmt2d*^*-/-*^ cells supports this new finding as a direct interaction between KMT2D and the *Shox2* promoter. We propose that loss of KMT2D (and thus H3K4me3) in chondrocytes and their precursors leads to decreased *Shox2* expression and subsequent release of *Sox9* inhibition; this leads to increased SOX9, which activates the chondrocyte differentiation program prematurely, including overexpression of *Col2a1* and *Col10a1*, and leads to precocious differentiation of chondrocytes.

Our finding of *Shox2* as a potential direct target of KMT2D is intriguing and quite relevant to the KS1 phenotype. A transcription factor highly conserved between mouse and humans (38,39), *Shox2* is expressed in developing limbs, the palate, the central nervous system, and the heart in both species (38), as well as in additional pharyngeal arch-derived craniofacial structures and the nasal process in humans. Notably, all of these tissues are affected in KS1 with its associated findings of developmental delay/intellectual disability, short stature/lower limb shortening, congenital heart disease, and well-documented craniofacial phenotype, including flattening of the facial profile due to short columella and depressed nasal tip, as well as ear and palate abnormalities and hearing loss (5,40). No congenital disorders in humans have yet been attributed to *SHOX2* intragenic mutations to determine whether this might lead to a phenocopy of KS. Contiguous gene deletions and duplications involving human chromosome 3q25-3q26 including *Shox2* and/or its known regulatory elements have been described, however, and these individuals have intellectual disability, skeletal and growth abnormalities, and dysmorphic facial features (41-43), resembling aspects of the KS phenotype. Neural-specific conditional ablation of *Shox2* leads to poor cerebellar development associated with precocious differentiation of neural progenitors and motor coordination deficits (44), supporting a functional role for SHOX2 in neurological development and potentially implicating it in motor delays, a key aspect of the KS1 phenotype. *Shox2* ^*-/-*^ mouse embryos die during mid-gestation and exhibit cleft palate and heart defects (45), features often seen in KS, though it is unclear whether additional skeletal growth phenotypes were examined. Conditional ablation of *Shox2* in mesenchymal cells and early chondrocytes leads to shortening of femurs and humeri (26), resembling the femur shortening observed in *Kmt2d*^*+/βGeo*^ mice. In their studies this was associated with precocious chondrocyte differentiation and hypertrophy, similar to what we observed in *Kmt2d*^*-/-*^ stable chondrocyte cell lines and at *Kmt2d*^*+/ βGeo*^ growth plates. Moreover, the conditional ablation in mesenchymal cells at an early time point is associated with increased expression of *Col2a1*, an early marker of chondrocyte differentiation, and *Sox9*, the master regulator of chondrogenesis (26), similar to our findings in *Kmt2d*^*-/-*^ and *Kmt2d*^*ΔR5551/-*^ stable chondrocyte cell lines. Finally, later conditional ablation in developing chondrocytes results in increased expression of late markers of chondrocyte differentiation like *Col10a1* (26), similar to our findings in *Kmt2d*^*-/-*^ cells. These results support our observations, and together the findings suggest a mechanism whereby loss of *Shox2* expression in chondrocytes or their precursors, whether genetic or epigenetic in nature, leads to *Sox9*-mediated precocious chondrocyte differentiation and ultimately to shortening of long bones.

Transient overexpression of *Shox2* in *Kmt2d*^*-/-*^ cells reduced *Sox9* transcript levels back toward those observed in *Kmt2d*^*+/+*^ cells. Although there was no significant difference between groups, *Sox9* levels in *Kmt2d*^*-/-*^ cells overexpressing *Shox2* did not appear to be reduced completely to levels observed in *Kmt2d*^*+/+*^ cells. The reduction in *Sox9* appears insufficient to restore proper expression levels of *Col2a1* (and *Col10a1*) and rescue the cellular phenotype, as evidence by the lack of reduction in alcian blue staining upon overexpression of *Shox2* in *Kmt2d*^*-/-*^ cells. While we observed decreased alcian blue staining in some images, this was not consistent. This discrepancy may be due to the transient nature of *Shox2* overexpression in our system and the extended length of time required to observe changes in alcian blue staining over the course of differentiation (14 days). Therefore, even a slight decrease in alcian blue staining may be suggestive of partial restoration of the cellular differentiation phenotype. Alternatively, there are many other target genes and pathways involved in KS-associated precocious chondrocyte differentiation. Indeed we observed thousands of differentially expressed genes upon differentiation of mesenchymal cells to chondrocytes and roughly a thousand differentially expressed genes when we compared *Kmt2d*^*-/-*^ and *Kmt2d*^*+/+*^ cells in both the differentiated and undifferentiated state. Restoration of one KMT2D target gene, even if critically important, simply may not be sufficient to rescue the cellular differentiation phenotype. Our findings here support the idea that multiple key cell type-specific target genes are disrupted and lead to relevant phenotypes in KS and other Mendelian disorders of the epigenetic machinery (1).

It is worth mentioning that *mShox2*, our novel KMT2D target, is the most closely related gene to the better-known *hSHOX* gene, which is known to cause multiple short stature syndromes with associated mesomelia (23-25) and to contribute to the short stature seen in Turner syndrome (23, 46). Moreover, a subset of individuals with Turner syndrome have a KS phenotype (47-50). *SHOX* and *SHOX2* are thought to exhibit functional redundancy in humans with no known mouse *SHOX* homologue (38, 39), suggesting that *SHOX2* may perform the overlapping functions of both genes in mice (39). In humans, *SHOX* is thought to mainly pattern the distal limb segment (tibia/radius) while *SHOX2* may be more important in patterning the proximal limb segment (femur/humerus; 39). Because both proximal femurs and distal tibias are shortened in *Kmt2d*^*+/βGeo*^ mice, our studies fit with prior work and suggest *SHOX2* may indeed serve dual functions. Based on our molecular studies in mice and data from humans with KS1, we hypothesize that *SHOX2* may be a target of KMT2D in humans as well, and we argue that *SHOX2* is a more likely target than *SHOX* based on two observations. First, a hallmark of every *SHOX-*associated disorder is mesomelic upper limb shortening, and humans with KS1 were shown to have disproportionately long arms (31), disfavoring *SHOX* as a key target of KMT2D and mediator of skeletal growth retardation in human KS1. Second, the expression patterns of *mShox2* and *hSHOX2* are more similar than *mShox2* and *SHOX*, and the tissue distribution of expression of *hSHOX2* overlaps almost exactly with the key tissues affected in KS1, including skeletal and others. Because *SHOX* is expressed in a subset of those tissues, we cannot rule out its involvement; however, our findings favor *SHOX2* as a mediator of key aspects of KS1.

Our findings reveal the first mechanistic insights into the molecular basis of the skeletal growth retardation phenotype and the highly characteristic facial appearance of KS1. We have characterized and quantified a robust skeletal growth retardation phenotype in KS1 mice, which resembles the human condition and involves disrupted endochondral ossification at distinct sites. Furthermore, we have elucidated a novel mechanism involving loss of KMT2D-mediated H3K4me3 at a new target (*Shox2*) and release of SOX9 inhibition, which allows precocious chondrocyte differentiation to proceed unchecked. Partial rescue of the KS1 chondrocyte gene expression profile through modulation of a single disrupted target gene, *Shox2*, fits with our model that KMT2D acts on multiple targets, some of which have key functional consequences. This is encouraging for future development of therapeutics directed against key epigenetic machinery targets implicated in KS1 and related disorders but suggests a broader therapeutic approach may be required in some cases. The previously identified roles of SHOX2 in skeletal tissues including long bones and the palate, as well as in other tissues relevant to KS, like cerebellum, point to additional potential roles for this gene in the pathogenesis of KS1 and related disorders and suggest additional therapeutic benefits to targeting *SHOX2* for the treatment of diverse manifestations of KS, other Mendelian disorders of the epigenetic machinery, and more common disease states that disrupt normal growth and development.

## Materials and Methods

### Mice

*Kmt2d*^*+/βGeo*^ mice have been previously described (15, 16). Mice were housed in a clean, specific pathogen-free state-of-the-art animal facility in ventilated racks and provided *ad lib* access to a standard rodent diet and to filtered water via an automatic watering system. All mice used in these studies were approximately 6 weeks of age unless otherwise noted, and littermates were used as controls in all experiments. Analyses were performed in a blinded manner unless otherwise noted. A single *Kmt2d*^*+/ +*^ mouse was excluded from all analyses due to being obviously runted and for having gross midfacial asymmetery related to incisor malocclusion, making this animal an outlier with respect to size, craniofacial shape/asymmetry, and micro-CT parameters. Otherwise, no outliers were excluded from any comparisons or statistical calculation.

### High-resolution micro-computed tomography

Femurs, tibias, and skulls were fixed in 4% paraformaldehyde, washed, and transferred to 70% ethanol. High-resolution images were acquired using a desktop micro-tomographic imaging system (Skyscan 1172, Bruker) in accordance with the recommendations of the American Society for Bone and Mineral Research (ASBMR;(51). Full length scans were reconstructed with NRecon software (Bruker). Femurs and tibias were scanned at 65 keV and 153μA using a 1.0 mm aluminum filter with an isotropic voxel size of 10 μm. In the femur, trabecular bone parameters were assessed in a region of interest 500 μm proximal to the growth plate and extending for 2 mm (200 CT slices) using CtAn software (Bruker). Cortical bone structure was assessed in the femur using a 500μm region of interest centered on the mid-diaphysis. Skulls were scanned at an isotropic voxel size of 10-28 μm at 80 keV and 120μA.

### Craniofacial morphometric analysis

We used geometric morphometrics to test the effect of the *Kmt2d*^*+/βGeo*^ mutation on craniofacial structure. MicroCT image volumes of the head were obtained for a comparative adult sample of *Kmt2d*^*+/βGeo*^ mice (N=13) and *Kmt2d*^*+/+*^ littermates (N=21), as described above (scan parameters: 80 keV and 120μA.; reconstruction: 0.010-0.028mm cubic voxels). Images were reconstructed using Amira post-processing software (v. 6.1.1, FEI), and 3D models of the skull were extracted based on density thresholds. The shape of the cranium was estimated by collecting three-dimensional coordinate data in Amira for biologically relevant, homologous landmarks (K=18; see **Fig. 1E**). We used MorphoJ software to produce a Procrustes superimposition of all landmark configurations (52) and, based on the assumption of object symmetry for the skull, analyzed only the symmetric portion of variance to minimize the effects of subtle asymmetries (53). Principal components analysis (PCA) was applied to visualize the most influential patterns of shape variance within the combined sample, and Procrustes ANOVA was used to test the effect of the mutation on overall cranial shape.

### Histology and histomorphometry

Tibias and skulls were fixed in 4% paraformaldehyde and then transferred to 10-14% EDTA for decalcification. Decalcified tibias and skulls were then processed, embedded, sectioned, and stained with hematoxylin and eosin (H and E). Images were taken using a Nikon 80i microscope and analyzed using NIS elements software. For tibias, longitudinal growth plate sections were used, and proliferative zone, hypertrophic zone, and total growth plate heights were measured in at least 3 sites per section in 4 sections per mouse within the central part of the growth plate. Number of cells per column within the proliferative zone and within the hypertrophic zone were also counted. For skulls, parasagittal sections were used, and intrasphenoidal synchondrosis growth plate heights were measured at 5 sites per section and in 3 sections per mouse within the central two-thirds of the growth plate. For all, an average measurement or cell count per mouse was generated and then the mean within each experimental group was calculated.

### Cell culture

ATDC5 cell line (19; Sigma-Aldrich; 99072806) was obtained from European Collection of Authenticated Cell Cultures. ATDC5 cells were maintained in standard medium: DMEM/Ham’s F12 medium containing 5% fetal bovine serum (FBS), antibiotics (100 units/ml penicillin, 100 μg/ml streptomycin), and 2 mM L-glutamine in 5% CO_2_ in a 37 °C incubator.

### Generation of stable cell lines

ATDC5 cells were seeded at 2 × 10^5^ cells/well in 6-well plates overnight. CRISPR-Cas9 constructs were a gift from J. Robertson and L. Goff. pSpCas9(BB)-2A-Puro (PX459) V2.0 vector containing gRNA inserts targeting *Kmt2d* exon 51 (5’-TCTGGCTCGTTCG CGTATCC-3’) and exon 53 (5’-TCCTTTGGGGATTCGCCGGC-3’) or empty vector were transfected using Lipofectamine 3000 (Invitrogen) according to the manufacturer’s instructions. 24 hrs post-transfection, the cells were treated with puromycin at a final concentration of 5 μg/mL for 3 days and the cells were allowed to recover. For single cell clonal analysis, cells were trypsinized and plated in 96 well plates at average 1 cell/well or plated in 100 mm cell culture dish at average 168 or 336 cells/dish and incubated at 37 °C for two weeks. Each well or plate was microscopically evaluated, and single cell-derived clones were selected, expanded, and genotyped using colony PCR and Sanger sequencing. PCR conditions available on request. Primers used were as follows: 5′-ACTCCAAGTCATCTCAGTAC-3′ and 5′-ACTGATAGTCATAGGTCAGC-3′. KMT2D protein expression was detected by Western blot (**Supplemental Figure 4**). *Kmt2d*^*-/-*^ stable cell lines have bi-allelic deletions within the catalytic SET domain, and *Kmt2d*^*ΔR5551/-*^ stable cell lines each have a monoallelic deletion within the catalytic SET domain on one allele and deletion of a single amino acid Arg5551 on the other allele. Arg5551 has not been associated with disease and corresponds to the site in the mRNA transcript expected to be cut by Cas9 based on the targeting strategy. Antibody specific for KMT2D (Millipore; ABE1867) was used to verify protein levels by Western blot analysis.

### Chondrocyte differentiation

ATDC5 stable cell lines (*Kmt2d*^*+/+*^, *Kmt2d*^*ΔR5551/-*^, *Kmt2d*^*-/-*^) were seeded in the above medium at 1× 10 ^5^ cells/well in 6-well plates (alcian blue staining; RNA isolation), at 1×10 ^3^ cells/well in 96-well plates (MTT assay), and at 6.3 × 10 ^3^ cells/well in a 24-well plates (cell counting). The next day and every 2-3 days thereafter, medium was replaced with chondrogenic differentiation medium: DMEM/ Ham’s F12 supplemented with 5% FBS, 100 units/ml penicillin, 100 μg/ml streptomycin, 2 mM L-glutamine, 1X Insulin-Transferrin-Selenium (Gibco), 50 μg/ml L-ascorbic acid and 10 mM β-glycerophosphate (MilliporeSigma). Cells were harvested at 0, 4, 7, 14, and 21 days.

### Alcian blue staining

At the indicated time points, cells were fixed in 4% paraformaldehyde, stained with alcian blue (Sigma-Aldrich), and microscopically evaluated. After permeabilization with 1% SDS, absorbance was measured at 605 nm using a Biotek synergy 2 plate reader.

### MTT cell proliferation assay

At the indicated time points, 10 μL of cell proliferation reagent (MTS) was added to the medium and incubated at 37° C, 5% CO_2_ for 1.5 hrs, as per manufacturer’s recommendation (Promega). Absorbance was measured at 490 nm using a Biotek synergy 2 plate reader.

### Cell counting assay

At the indicated time points, cells were trypsinized and stained with a 0.4 % trypan blue solution (Corning). Cells were manually counted using a hemocytometer.

### Quantitative RT-PCR

Total RNA was isolated using TRIZOL reagent according to the manufacturer’s instructions (Invitrogen) at the indicated time points. Reverse transcription was performed using the SuperScript IV First-Strand Synthesis System (ThermoFisher Scientific). Quantitative real-time PCR analysis using the comparative Ct method was performed on the Applied Biosystem Vii 7 system (ThermoFisher Scientific) with the Powerup SYBR Master Mix (ThermoFisher Scientific) according to the manufacturer’s instructions.

### Chromatin immunoprecipitation

ChIP assay was performed according to Cold Spring Harbor protocol (54). Briefly, cells were cross-linked for 10 min with formaldehyde at a 1% final concentration, and the reaction was quenched by adding glycine to a final concentration of 137.5 mM. The cells were then rinsed in ice-cold PBS, scraped, and resuspended in a lysis buffer with the addition of complete protease inhibitor cocktail (Cell Signaling Technology). Chromatin was sheared by sonication 3×5 cycles of 30 sec ON/30 sec OFF with high intensity (Bioruptor) to generate DNA fragments between 300-500 bp, used for immunoprecipitation with anti-H3K4me3 (Millipore Sigma), and then captured by protein G-agarose/salmon sperm DNA (Millipore Sigma). Precipitated DNA was reverse cross-linked and then amplified by qPCR using primers amplifying amplicons with the following midpoints relative to the transcription start site of *Shox2:* −500, −250, 0 (TSS), +200, and +525 and compared with the amount of input DNA before immunoprecipitation.

### RNA-seq

ATDC5 stable cell lines (*Kmt2d*^*+/+*^, *Kmt2d*^*ΔR5551/-*^, *Kmt2d*^*-/-*^) were seeded at 1× 10 ^5^ cells/well in 6-well plates and allowed to adhere overnight in standard medium. The next day and every 2-3 days thereafter, medium was replaced with chondrogenic differentiation medium. Total RNA was isolated using TRIZOL reagent and RNA Clean & Concentrator-5 kit (Zymo Research) according to the manufacturers’ instructions. Contaminating genomic DNA was removed by treatment with DNase-I. RNA was assessed for quantity and quality using Qubit RNA BR Assay Kit (ThermoFisher Scientific) and RNA 6000 Nano Kit on the Agilent Bioanalyzer 2100 system (Agilent Technologies), respectively, according to the manufacturers’ instructions. mRNA was purified from 1 μg of total RNA (≥100 ng/μL) using NEBNext Poly(A) mRNA Magnetic Isolation Module for Illumina (New England Biolabs). Sequencing libraries were generated using NEBNext Ultra II RNA Library Prep with Sample Purification Beads for Illumina (New England Biolabs) in accordance with the manufacturer’s recommendations and were validated using the Agilent High Sensitivity DNA assay on the Agilent Bioanalyzer 2100 system and quantified by NEBNext Library Quant Kit for Illumina (New England Biolabs). After clustering of the index-coded samples, libraries were sequenced on an Illumina HiSeq 2500 platform which generated 100 bp single-end reads. RNA-seq was performed on two (*Kmt2d*^*-/-*^) or three (*Kmt2d*^*+/+*^) biological replicates for each differentiation state (chondrocytes vs undifferentiated cells), and each biological replicate is a distinct clonal cell line. Two technical replicates were performed for each cell line.

### RNA-seq bioinformatics analysis

We first pseudoaligned the reads to a fasta file (Mus_musculus.GRCm38.cdna.all.fa.gz) obtained from Ensembl (http://uswest.ensembl.org/Mus_musculus/Info/Index, version 91, downloaded January 2018), which contained all mouse cDNA sequences, and then performed the quantification of transcript abundances using Salmon (55). Then, we utilized the tximport R package (56) in order to obtain normalized gene-level counts from the transcript abundances. To achieve this, we set the “countsFromAbundance” parameter equal to “lengthScaledTPM”. Subsequently, using the edgeR (57) and limma (58) R packages, we applied a log_2_ transformation to the gene-level counts and normalized each sample with the “voom” function in limma, using the effective library size (i.e. the product of the library size and the normalization factors, calculated using the “calcNormFactors” function in edgeR). We then estimated the mean-variance relationship and computed weights for each observation. Since the differential expression analysis included technical replicates for each of three wild-type and two mutant clones, we fit a mixed linear model using the function “duplicateCorrelation” from the statmod R package (60), by blocking on the clone to account for the correlation among technical replicates. Finally, we performed the differential analysis with the limma R package with an FDR of 0.05 as the threshold for statistical significance. Prior to performing the principal components analysis, we converted transcript abundances to gene-level counts using the tximport R package by setting the “countsFromAbundance” parameter to “no”. Then, we first used the “vst” function from the DESeq2 R package (59); with the parameter “blind” set to “TRUE”) in order to apply a variance stabilizing transformation to the obtained gene-level counts. Then, without standardizing the resulting expression matrix, we used the 1000 most variable genes to estimate the principal components.

### Lentiviral transduction

Lenti-ORF particles, Shox2 (Myc-DDK-tagged) and Control (Myc-DDK-tagged) were purchased from Origene. To transduce the ATDC5 cells, 1.2 × 10 ^5^ cells/well were seeded in 6-well plates overnight and then transduced with the lentivirus at a multiplicity of infection (MOI) of 5 and with a final polybrene concentration of 8 ug/mL.

### Statistics

One-sided unpaired student’s t-test was used to calculate p values, unless otherwise noted; p < 0.05 indicated statistical significance.

### Study approval

All experiments using laboratory mice were performed in accordance with the NIH Guide for the Care and Use of Laboratory Animals and were approved by the Johns Hopkins Animal Care and Use Committee and performed in accordance with their guidelines.

## Author contributions

JAF, HTB designed, initiated, oversaw, and directed the study and wrote the manuscript; JAF, RCR, WYL, SC, SEL, TL performed experiments and acquired data; JAF, WYL, RCR, VDL, LB, KH analyzed data.

## Acknowledgements

We thank Dr. Joel Benjamin, Dr. Jefferson Doyle, and Dr. Genay Pilarowski for their work in the early stages of this project. We thank Dr. Li Zhang for technical advice throughout the project. We thank Dr. Thomas Clemens for many helpful suggestions throughout the study and for reading the manuscript, Dr. Rosa Serra and George Coricor for their advice regarding the use of ATDC5 cells, Dr. Timothy Cox for advice on skull histology, and Dr. Janet Crane for advice on long bone histology. We thank Dr. Loyal Goff and Johanna Robertson for sharing *Kmt2d* targeting constructs. We thank Dr. Barbara Migeon for critical reading of the manuscript. We thank Catherine Kiefe for assistance with creating the highlighter schematic. This study makes use of data generated by the DECIPHER community. A full list of centres who contributed to the generation of the data is available from http://decipher.sanger.ac.uk and via email from. Funding for DECIPHER was provided by the Wellcome Trust. The research in this manuscript was supported by a Baltimore Center for Musculoskeletal Science 2015 Pilot and Feasibility Award (J.A.F.); a grant from the William and Ella Owens Medical Research Foundation (J.A.F.); a Johns Hopkins School of Medicine Clinician Scientist Award (J.A.F.); a Hartwell Foundation Individual Biomedical Research Award (J.A.F.); National Institutes of Health grants to J.A.F. (1K08HD086250-01A1) and H.T.B. (DP5OD017877); and a grant from the Louma G. Foundation (H.T.B.).

