## Supplemental Figures SF1-SF8 for "Precocious Chondrocyte Differentiation Disrupts Skeletal Growth in Kabuki Syndrome Mice"

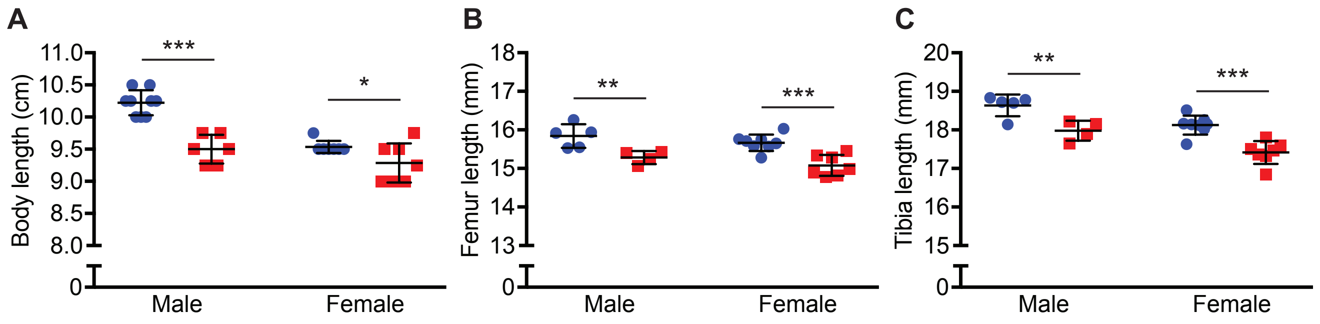


**Supplemental Figure 1: Kabuki syndrome 1 mice have consistent skeletal growth retardation at 18 weeks. (A)** Quantification of body length in 18 week old *Kmt2d*$^{+/+}$ male (n=9) and female (n=7) mice and *Kmt2d*$^{+/\beta Geo}$ male (n=6) and female (n=7) mice. Quantification of femur **(B)** and tibia **(C)** length in 18 week old *Kmt2d*$^{+/+}$ male (n=5) and female (n=8) mice and *Kmt2d*$^{+/\beta Geo}$ male (n=4) and female (n=7) mice. *Kmt2d*$^{+/+}$ mice represented by blue circles; *Kmt2d*$^{+/\beta Geo}$ mice represented by red squares. Data represent mean +/- standard deviation. *p<0.05, **p<0.01, ***p<0.001.

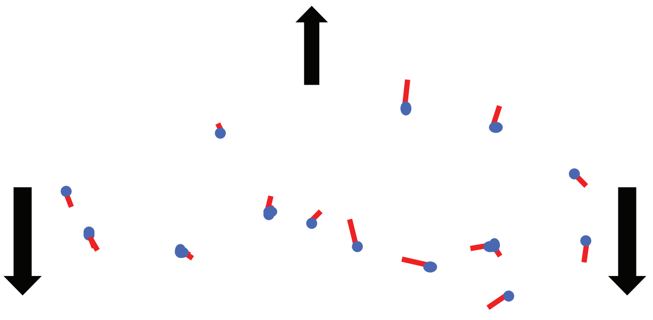


**Supplemental Figure 2: *Kmt2d*$^{+/\beta Geo}$ mice exhibit ventral bowing, brachycephaly, and dorsal expansion of the skull compared to *Kmt2d*$^{+/+}$ mice.** Blue points represent landmarks and show mean form of the sample in left lateral view. Red vectors show displacement of landmarks from the mean form in *Kmt2d*$^{+/\beta Geo}$ mice. Thick black arrows illustrate overall direction of displacement, suggesting ventral bowing, dorsal expansion, and brachycephaly. n=21 *Kmt2d*$^{+/+}$ mice, n=13 *Kmt2d*$^{\beta geo/+}$ mice.

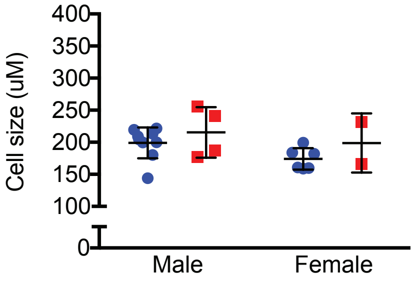


**Supplemental Figure 3:** ***Kmt2d***$^{+/\beta Geo}$ **mice exhibit no difference in hypertrophic chondrocyte cell size compared to** ***Kmt2d***$^{+/+}$ **mice.** Hypertrophic chondrocytes from H and E stained proximal tibia growth plate sections were measured and analyzed for cell size using Image J. Blue circles represent *Kmt2d*$^{+/+}$ mice (n=15; 9 males and 6 females); *Kmt2d*$^{+/\beta Geo}$ mice represented by red squares (n=6; 4 males and 2 females). Data represent mean +/- standard deviation.

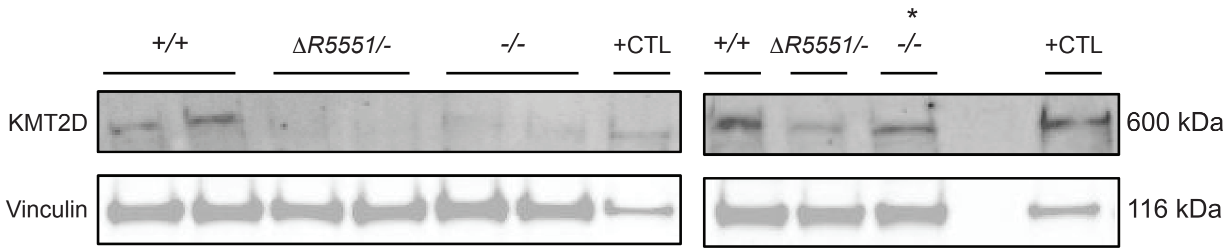


**Supplemental Figure 4: KMT2D protein expression in *Kmt2d*$^{+/+}$*, Kmt2d*$^{\Delta R5551/-}$*,* and *Kmt2d*$^{-/-}$ stable cell lines.** *Kmt2d*$^{+/+}$ stable cell lines express KMT2D (600kDa), similar to the positive control cell lysate (+CTL). *Kmt2d*$^{\Delta R5551/-}$ and *Kmt2d*$^{-/-}$ express little to no KMT2D protein, suggesting a hypomorphic and/or loss of function mutation mechanism. The *Kmt2d*$^{-/-}$ clonal cell line indicated with an asterisk continues to express KMT2D protein at similar levels to *Kmt2d*$^{+/+}$ stable cell lines and therefore was not used in experimental analyses. Vinculin was used as loading control.

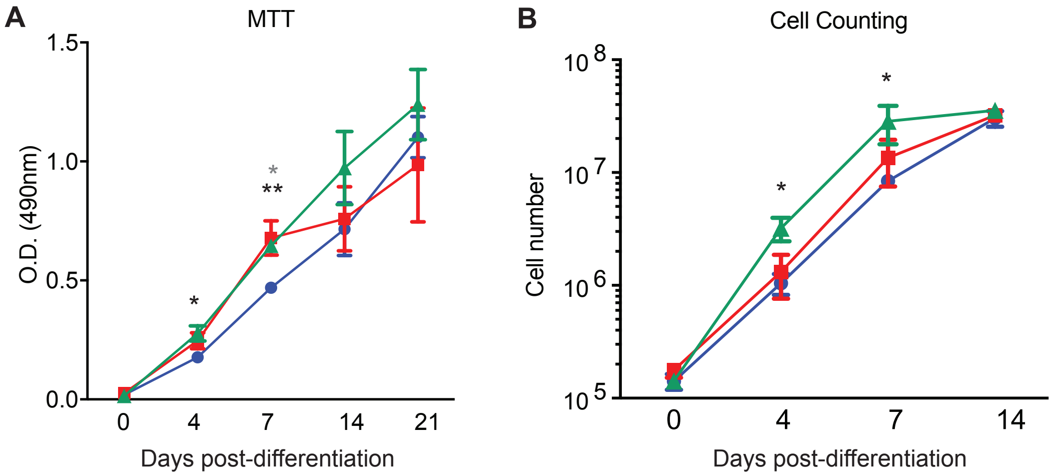


**Supplemental Figure 5. Cell proliferation is increased in *Kmt2d*$^{-/-}$ chondrocytes compared to *Kmt2d*$^{+/+}$ chondrocytes at days 4 and 7 post-induction of differentiation.** Cellular proliferation was measured using the MTT assay **(A)** and cell counting **(B)** both prior to differentiation and at 4, 7, and 14 days after induction of differentiation. Blue circles represent *Kmt2d*$^{+/+}$ cells, red squares represent *Kmt2d*$^{\Delta R5551/-}$ cells, and green triangles represent *Kmt2d*$^{-/-}$ cells. Error bars indicate standard error of the mean**.** Black asterisks represent differences between *Kmt2d*$^{+/+}$ and *Kmt2d*$^{-/-}$ cells while the gray asterisk represents differences between *Kmt2d*$^{+/+}$ and *Kmt2d*$^{\Delta R5551/-}$. *p value <0.05; **p value <0.01.

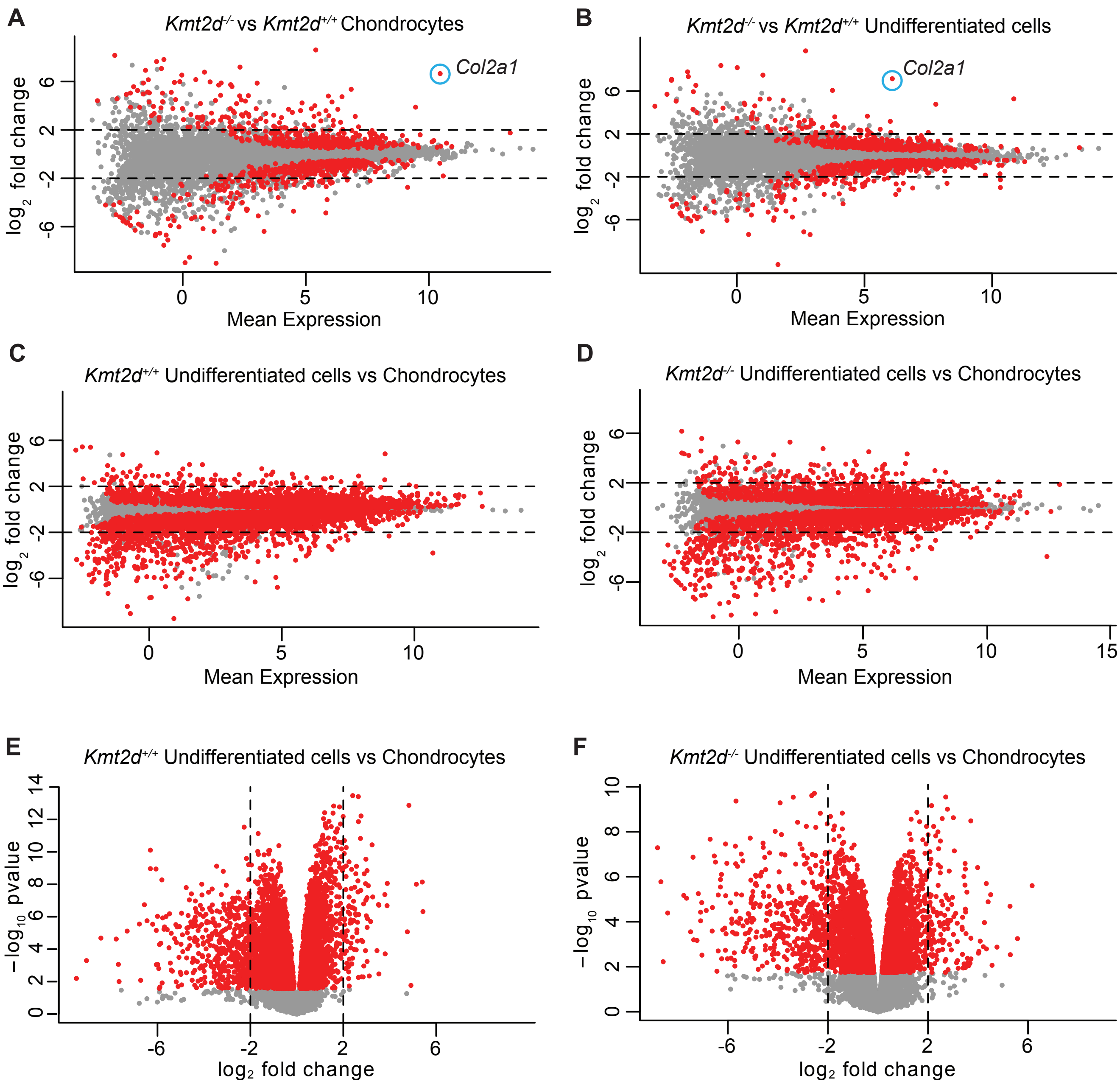


**Supplemental Figure 6: The number of genes required to undergo chondrocyte differentiation greatly exceeds the number altered upon loss of *Kmt2d*.** MA plots comparing differentially expressed genes in *Kmt2d*$^{-/-}$ and *Kmt2d*$^{+/+}$ chondrocytes **(A)** and undifferentiated mesenchymal cells **(B).** MA plots **(C, D)** and volcano plots **(E, F)** comparing differentially expressed genes in *Kmt2d*$^{+/+}$ **(C, E)** and *Kmt2d*$^{-/-}$ **(D, F)** cells over the course of differentiation from mesenchymal cells to chondrocytes.

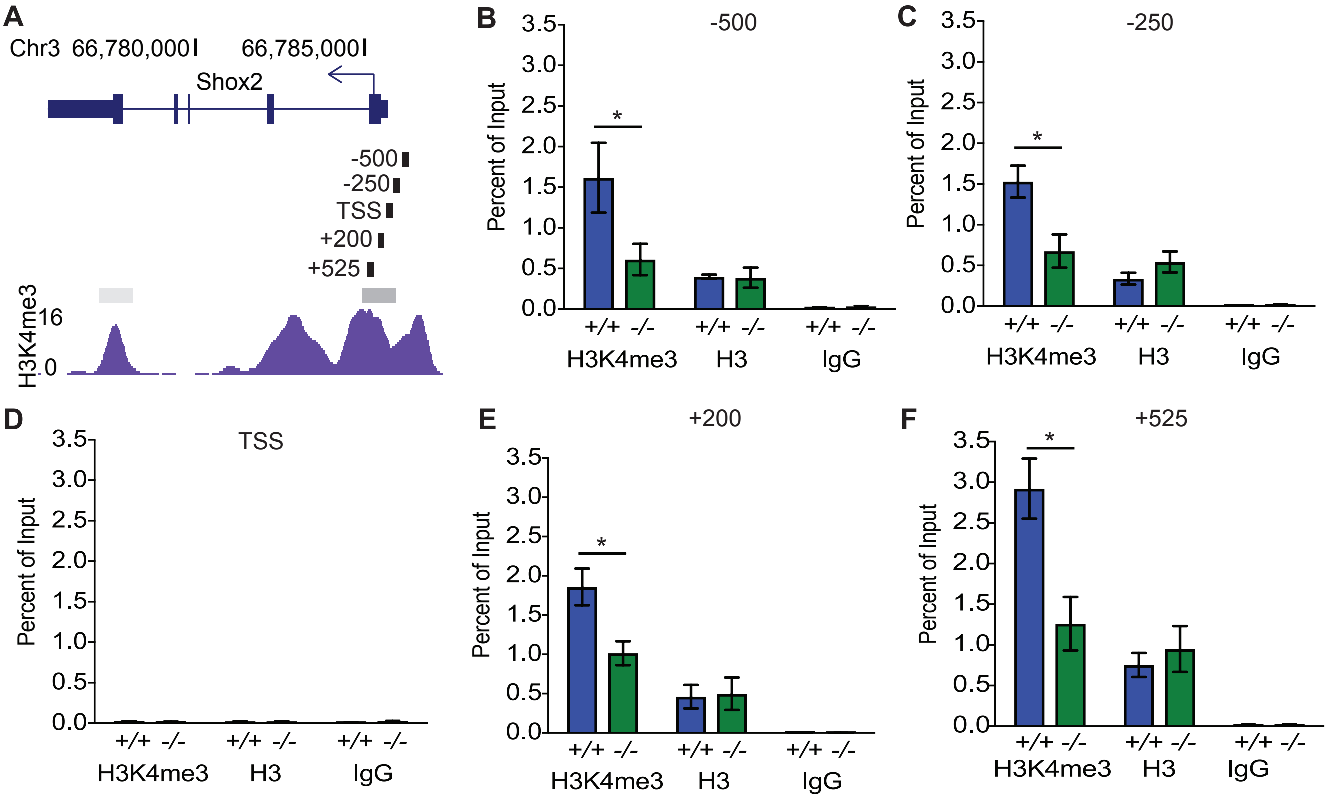


**Supplemental Figure 7. H3K4me3 levels are depleted surrounding the *Shox2* transcription start site in *Kmt2d*$^{-/-}$ cells whereas levels of H3 remain constant. (A)** Schematic indicating the genomic location of the primers used for quantitative PCR following chromatin immunoprecipitation (ChIP-qPCR) in relation to the levels and known peak of H3K4me3 at the 5' end of the *Shox2* gene in mouse embryonic limb bud from the ENCODE data set. The *Shox2* gene and its orientation, exons, and 5' and 3' untranslated regions are indicated with genomic coordinates above. Black vertical bars indicate positions of amplicons amplified by primer sets, and the adjacent numbers indicate the approximate midpoint of each amplicon in relation to the transcription start site (TSS) of *Shox2.* Solid gray horizontal bars indicate peaks of H3K4me3 with the darker gray bar indicating the peak of interest. The purple plot indicates relative H3K4me3 enrichment. ChIP-qPCR revealed depletion of H3K4me3 in *Kmt2d*$^{-/-}$ cells compared to *Kmt2d*$^{+/+}$ cells both upstream **(B, C)** and downstream **(E, F)** of the transcription start site. Levels of H3 remained constant in *Kmt2d*$^{-/-}$ and *Kmt2d*$^{+/+}$ cells upstream **(B, C)** and downstream **(E, F)** of the transcription start site. Neither H3K4me3 nor H3 was found at the transcription start site **(D)** in the presumed nucleosome-free region. Anti-IgG antibody was used as a negative control. Blue bars represent *Kmt2d*$^{+/+}$ cells*;* green bars represent *Kmt2d*$^{-/-}$ cells. Error bars indicate standard error of the mean**.** *p value <0.05.

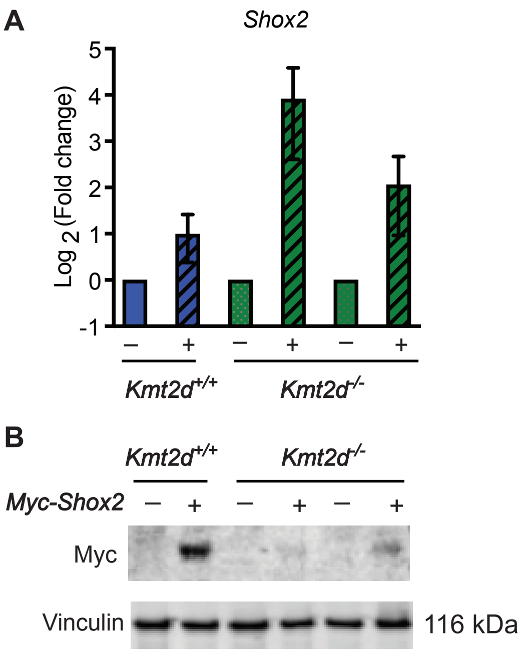


**Supplemental Figure 8. Increased *Shox2* transcript levels and Myc-SHOX2 protein overexpression in individual *Kmt2d*$^{-/-}$ and *Kmt2d*$^{+/+}$ stable cell lines.** *Shox2* overexpression by qRT-PCR with primers specific to *mShox2* **(A).** Blue bars represent *Kmt2d*$^{+/+}$ stable cell lines and green bars represent *Kmt2d*$^{-/-}$ stable cell lines. Solid bars represent cells infected with control lentivirus, and hatched bars represent cells infected with lentivirus overexpressing *Shox2.* **(B)** Western blot using anti-Myc antibody reveals Myc-SHOX2 overexpression in *Kmt2d*$^{+/+}$ cells and in one *Kmt2d*$^{-/-}$ cell line. Vinculin serves as a loading control.
